# The RAG1 Ubiquitin Ligase Domain Enhances the Assembly and Selection of T Cell Receptor Genes to Restrain the Autoimmune Hazard of Generating T Cell Receptor Diversity

**DOI:** 10.1101/2021.01.04.425211

**Authors:** Thomas N. Burn, Charline Miot, Scott M. Gordon, Erica J. Culberson, Tamir Diamond, Portia A. Kreiger, Katharina E. Hayer, Anamika Bhattacharyya, Jessica M. Jones, Craig H. Bassing, Edward M. Behrens

**Author notes:** Co-corresponding Authors: Craig H. Bassing, Ph.D. Children’s Hospital of Philadelphia, 4054 Colket Translational Research Building 3501 Civic Center Blvd., Philadelphia, PA 19104, Edward M. Behrens, M.D. Children’s Hospital of Philadelphia 1102 Abramson Research Center 3615 Civic Center Boulevard Philadelphia, PA 19104. Co-first Authors (Thomas N. Burn, Charline Miot).

## Abstract

RAG1/RAG2 (RAG) endonuclease-mediated assembly of diverse lymphocyte antigen receptor genes by V(D)J recombination is critical for the development and immune function of T and B cells. However, this process creates highly self-reactive cells that must be properly selected to suppress autoimmunity. The RAG1 protein contains a ubiquitin ligase domain that stabilizes RAG1 and stimulates RAG endonuclease activity *in vitro*. We report that mice with a mutation that inactivates the RAG1 ubiquitin ligase *in vitro* exhibit modestly reduced thymic cellularity, decreased assembly and altered repertoires of T cell receptor (TCR) β and α genes in thymocytes, and impaired thymocyte developmental transitions that require the assembly of TCRβ or α genes and signaling by their proteins. These RAG1 mutant mice also exhibit less efficient positive selection and superantigen-mediated negative selection of conventional αβ T cells, 2) impaired differentiation of iNKT lineage αβ T cells, and 3) CD4^+^ αβ T cells with elevated autoimmune potential. Our findings demonstrate that the RAG1 ubiquitin ligase domain functions *in vivo* to stimulate the assembly and selection of TCRβ and TCRα genes, thereby establishing replete diversity of αβ TCRs and αβ T cell lineages while restraining the inherent autoimmune hazard of generating diverse antigen specificities.

## Introduction

The ability of T and B lymphocyte populations to express antigen receptors able to recognize a potentially unlimited number of distinct pathogens is the fundamental basis of adaptive immunity. RAG1/RAG2 (RAG) endonuclease-mediated assembly of T cell receptor (TCR) and immunoglobulin (Ig) genes through V(D)J recombination establishes this vast pool of diverse receptors. Germline TCR and Ig loci consist of variable (V), joining (J), and sometimes diversity (D), gene segments residing upstream of constant (C) region exons. During T and B cell development, the lymphocyte-specific RAG complex cooperates with lineage- and developmental stage-specific activation of TCR and Ig loci to perform V(D)J recombination (1). RAG induces DNA double strand breaks (DSBs) adjacent to two participating gene segments and cooperates with DSB repair factors to create V(D)J coding joins that comprise the second exons of assembled V(D)J-C antigen receptor genes (1, 2). The broad utilization of high numbers of gene segments and imprecision in V(D)J coding join formation creating a hypervariable complementatry-determineing region (the CDR3 region) cooperate to produce a vast population of diverse antigen receptor genes. Due to imprecise means of opening and processing coding ends, the outcomes of V(D)J recombination include out-of-frame genes unable to make protein and receptors that recognize self-antigens and have potential for autoimmunity. Accordingly, developing T and B lymphocytes engage quality control checkpoints to select for potentially beneficial antigen receptor proteins and against hazardous highly self-reactive receptors.

Both immature T and B cells employ developmental programs that link functional protein expression from in-frame V(D)J rearrangements to signal either survival and continued differentiation or apoptosis (1). The maturation of αβ T cells in the thymus provides a paradigm for how V(D)J recombination and antigen receptor protein quality control checkpoints cooperate to create a large pool of lymphocytes with diverse receptors that provides immunity from pathogens without causing overt autoimmunity. After entering the thymus, early thymocyte progenitor cells differentiate into CD44^+^CD25^+^ stage 2 and then into CD44^-^ CD25^+^ stage 3 CD4^-^CD8^-^ double negative (DN) thymocytes (3, 4). Both DN2 and DN3 cells express RAG and conduct V(D)J recombination of *Tcrb*, *Tcrd*, and/or *Tcrg* loci (3, 4). Although DN2-to-DN3 thymocyte maturation can occur independent of RAG and V(D)J recombination, an in-frame *Tcrb* rearrangement is necessary for development beyond the DN3 stage (3, 4). The resultant TCRβ protein associates with invariant pTα protein to produce pre-TCR complexes that signal antigen-independent survival, proliferation, and differentiation of DN3 cells (β-selection) into CD44^-^CD25^-^ stage 4 DN thymocytes and then CD4^+^CD8^+^ double-positive (DP) thymocytes (3, 4). DP cells express RAG and conduct V(D)J recombination of *Tcra* loci, which typically proceeds through successive rounds of V-to-J rearrangement on both alleles (5). After an in-frame VJ rearrangement, the resulting TCRα protein can pair with TCRβ protein and yield surface αβ TCRs, which must signal to prevent death by neglect (6–9). This signal activation requires appropriate physical interactions between αβ TCRs and self-peptide/MHC (pMHC) complexes displayed on thymic epithelial cells (TECs) or dendritic cells (DCs), meaning that a functional αβ TCR is inherently self-reactive (6–9). Interactions below a particular low threshold of affinity (or avidity) cannot activate signalling to block apoptosis (6–9). At the other extreme, contacts above a higher threshold trigger strong signals that causes apoptosis (negative selection) to delete highly self-reactive cells with substantial autoimmune potential (6–9). Contacts between these thresholds activate TCR signals of a strength within a range that promotes survival and differentiation of DP cells (positive selection) into CD4^+^ or CD8^+^ single-positive (SP) thymocytes, which lack RAG expression and exit the thymus as mature αβ T cells (6–9). Strong TCR signals in DP cells can also activate an alternative agonist selection process that promotes differentiation of unconventional αβ T cell lineages: regulatory T (T reg) cells, intestinal intraepithelial lymphocytes (IELs), and invariant natural killer T (iNKT) cells (6, 7).

The RAG1 protein N-terminus has a Zinc-binding RING finger ubiquitin ligase domain of undetermined relevance for antigen receptor gene assembly. *In vitro,* RAG1 autoubiquitylates on lysine 233 (10), which enhances RAG endonuclease activity *in vitro* as evidenced by impaired cleavage upon mutation of this residue (11). *In vitro,* RAG1 also ubiquitylates other proteins including histone H3 within chromatin (12, 13). RAG1 RING domain mutations that disrupt ubiquitin ligase activity impair RAG-mediated cleavage and joining of chromatin substrates in cell lines, which could be due to loss of histone H3 ubiquitylation and/or alteration in RAG1 protein structure (13–15). The C328Y mutation of a critical structural zinc- binding cysteine of human RAG1 impairs V(D)J recombination and causes severe T and B cell immunodeficiency associated with uncontrolled proliferation of activated oligoclonal αβ T cells (16). The analogous C325Y mutation of mouse Rag1 protein (C325Y) *in vitro* destabilizes Rag1 tertiary structure, abrogates ubiquitin ligase activity, and reduces V(D)J recombination (22). Mice with homozygous Rag1 C325Y mutation have a near complete block of αβ T cell development at the DN3 thymocyte stage due to impaired *Tcrb* recombination (14, 15). RING domains have a conserved proline (proline 326 in mouse Rag1) that is critical for ubiquitin ligase activity by permitting functional interaction with ubiquitin- conjugating enzymes (17, 18). Relative to C325Y mutation, P326G mutation abrogates Rag1 ubiquitin ligase activity equivalently, but has substantially less severe effects on both destabilizing tertiary structure of the Rag1 RING domain and reducing RAG endonuclease activity (15). Accordingly, P326 mutation of endogenous mouse Rag1 protein allows an approach to elucidate potential physiologic roles of the RAG1 ubiquitin ligase domain beyond promoting V(D)J recombination by stabilizing RAG1 protein structure.

To determine potential *in vivo* functions of RAG1 ubiquitin ligase activity, we created and analyzed mice carrying a homozygous Rag1 P326G mutation. These mice exhibit phenotypes indicative of lower than normal RAG endonuclease activity including impaired B cell development, elevated Igκ^+^:Igλ^+^ B cell ratio, and decreased assembly of *Igh*, *Igκ*, *Igλ*, and *Tcrb* genes (19). We show here that these mice exhibit decreased levels and altered repertoires of *Tcrb* or *Tcra* rearrangements in DN3 or DP thymocytes, respectively, correlating with lower fractions of each cells selected for further differentiation. We find that the αβ TCR-signaled upregulation of CD69 is diminished during the initiation of positive selection. When assaying αβ TCRs expressing specific TCRβ or TCRα chains, we observe diminished efficiencies for superantigen-mediated negative selection of conventional αβ T cells and differentiation of iNKT lineage αβ T cells. Finally, we show that mature CD4^+^ effector αβ T cells of these mice exhibit normal immune responses to activation, yet possess greater intrinsic potential for autoimmunity. Our data demonstrate that the RAG1 ubiquitin ligase domain functions *in vivo* to stimulate TCRβ and TCRα gene assembly and αβ TCR selection, thereby generating replete αβ TCR diversity and αβ T cell lineages while restraining the inherent autoimmune hazard of establishing diverse TCR specificities. We discuss how the RAG1 ubiquitin ligase domain might function *in vivo* to regulate the assembly of TCRβ and TCRα genes, selection of αβ TCRs, and differentiation of αβ T cell lineages.

## Results

### Homozygous Rag1 P326G mutation diminishes TCR gene assembly and subsequent thymocyte developmental stage transitions

To determine potential *in vivo* functions of the RAG1 ubiquitin ligase, we created and analyzed C57BL/6 strain mice homozygous for the Rag1 P326G mutation (*Rag1^P326G/P326G^* mice; see Materials and Methods) that abrogates RAG1 ubiquitin ligase activity but minimally destabilizes RAG1 protein (15). We initially analyzed αβ T cell development and TCR gene rearrangements in *Rag1^P326G/P326G^* (*PG*) mice and wild-type (*WT*) mice. As compared to *WT* mice, *PG* mice have a ∼2-fold higher percentage and number of DN3 thymocytes and correspondingly lower percentage and number of DN4 thymocytes (Fig. 1, A and B). *PG* mice also exhibit ∼10% reductions in the percentage and number of DP thymocytes and the ratio of CD4^+^ SP thymocytes to DP thymocytes (Fig. 1, C-E). Despite these minor impairments of the DN3-to-DN4 and DP-to-SP thymocyte developmental transitions, *PG* mice have normal numbers of total thymocytes and splenic αβ T cells (Fig. 1 F). Using Taqman PCR to amplify Vβ rearrangements to Dβ1Jβ1.1, Dβ1Jβ2.1, or Dβ2Jβ2.1 complexes in non-selected DN3 cells, we observed ∼50% lower levels of rearrangement for nearly all individual Vβ segments in *PG* mice (Fig. 2, A and B). Notably, the recombination of some Vβ segments involving Jβ2.1 were undetectable in *PG* mice (Fig. 2, A and B), indicating altered Jβ repertoire from Rag1^P326G^ mutation. *Rag1* or *Rag2* mutations that substantially decrease RAG endonuclease activity, thymocyte development, and thymic cellularity, also alter the usage of individual Vβ segments (20, 21). However, high-throughput sequencing of *Tcrb* rearrangements in DN3 thymocytes shows similar usage of each Vβ segment in *PG* and *WT* mice (Fig. 2 C), reflecting the modest reduction of *Tcrb* gene assembly in *PG* mice. This analysis also reveals modest alterations in Dβ and Jβ usage, but no difference in CDR3β lengths in the *Tcrb* rearrangements of *PG* mice (Fig. 2, D and E; data not shown). Using Taqman PCR to amplify a subset of possible Vα-to-Jα rearrangements in non-selected DP thymocytes, we detected lower levels of rearrangements for many of the Vα/Jα combinations, particularly those involving the two most distal Jα gene segments (Jα17, Jα2) tested, in *PG* mice (Fig. 2, F). Together, these data indicate that homozygous Rag1^P326G^ mutation produces RAG endonuclease complexes that catalyze decreased levels of *Tcrb* and *Tcra* rearrangements, alter the primary repertoires of *Tcrb* and *Tcra* genes, and support less efficient thymocyte developmental transitions that depend on the functional assembly of a *Tcrb* or *Tcra* gene. The lower efficiency of overall *Tcrb* and *Tcra* gene assembly certainly contributes to the diminished DN3-to-DP and DP-to-SP thymocyte developmental progressions observed in *PG* mice.

**Figure 1.**
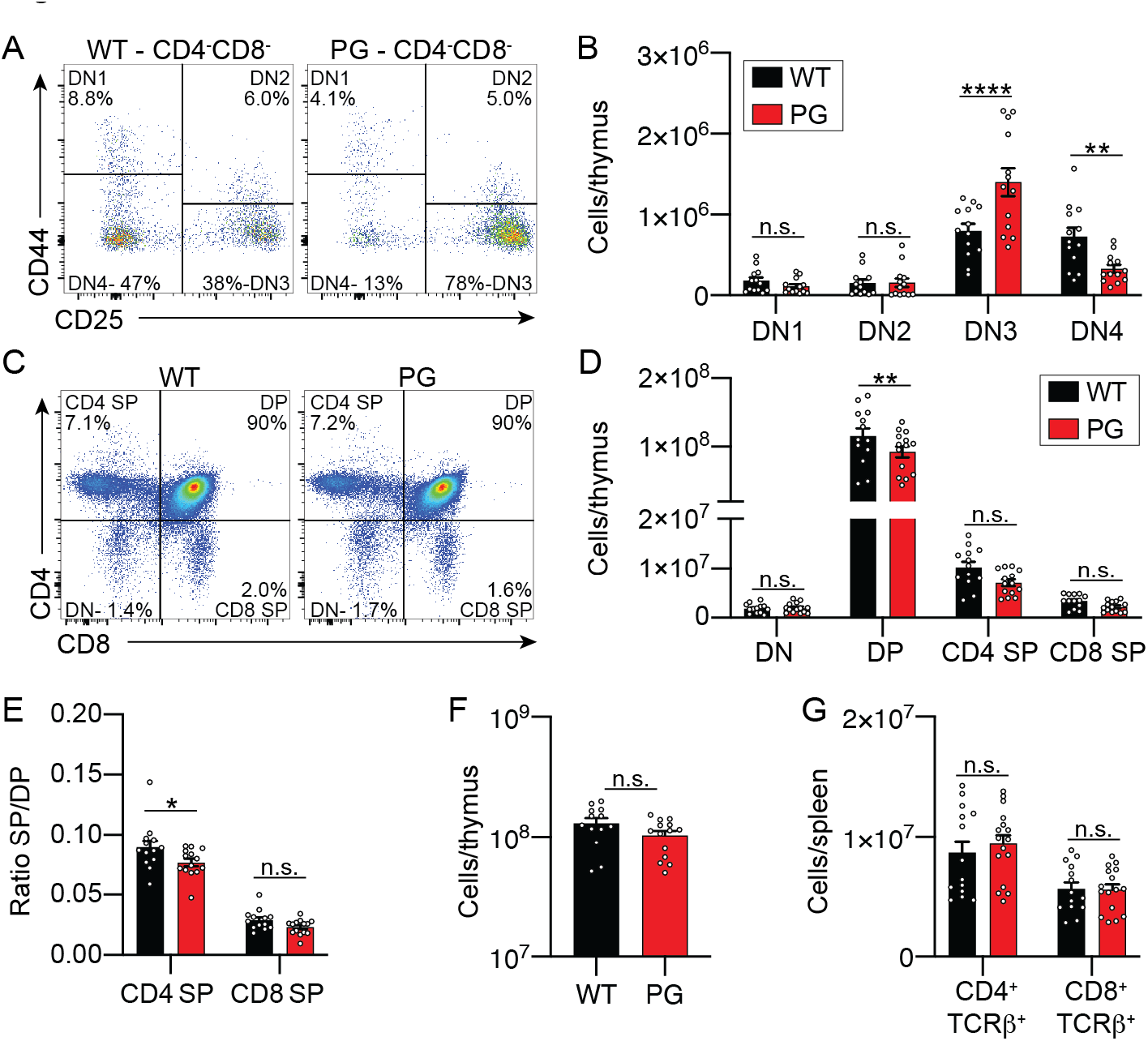
*PG* mice exhibit impaired thymocyte developmental transitions that require TCR gene assembly. **(A)** Representative flow cytometry plots of DN stages of thymocyte development showing the frequency of cells at each stage. **(B)** Quantification of numbers of cells in each DN stage. **(C)** Representative flow cytometry plots of DN, DP, CD4^+^ SP, and CD8^+^ SP thymocytes. **(D)** Quantification of numbers of cells in each stage. **(E)** Quantification of the ratio of CD4^+^ SP or CD8^+^ SP thymocytes to DP thymocytes. **(F-G)** Quantification of the numbers of total thymocytes (F) and splenic αβ T cells (G). (A-G) All data were collected from 6-10 weeks old mice and (B, D, E, F, and G) are combined from four independent experiments including a total of 14-16 mice per genotype. Bars indicate mean +/- SEM. Stats: 2-way ANOVA with Sidak multiple comparison test. n.s. p>0.05; * p<0.05, **p<0.01, *** p<0.001, **** p<0.0001.

**Figure 2.**
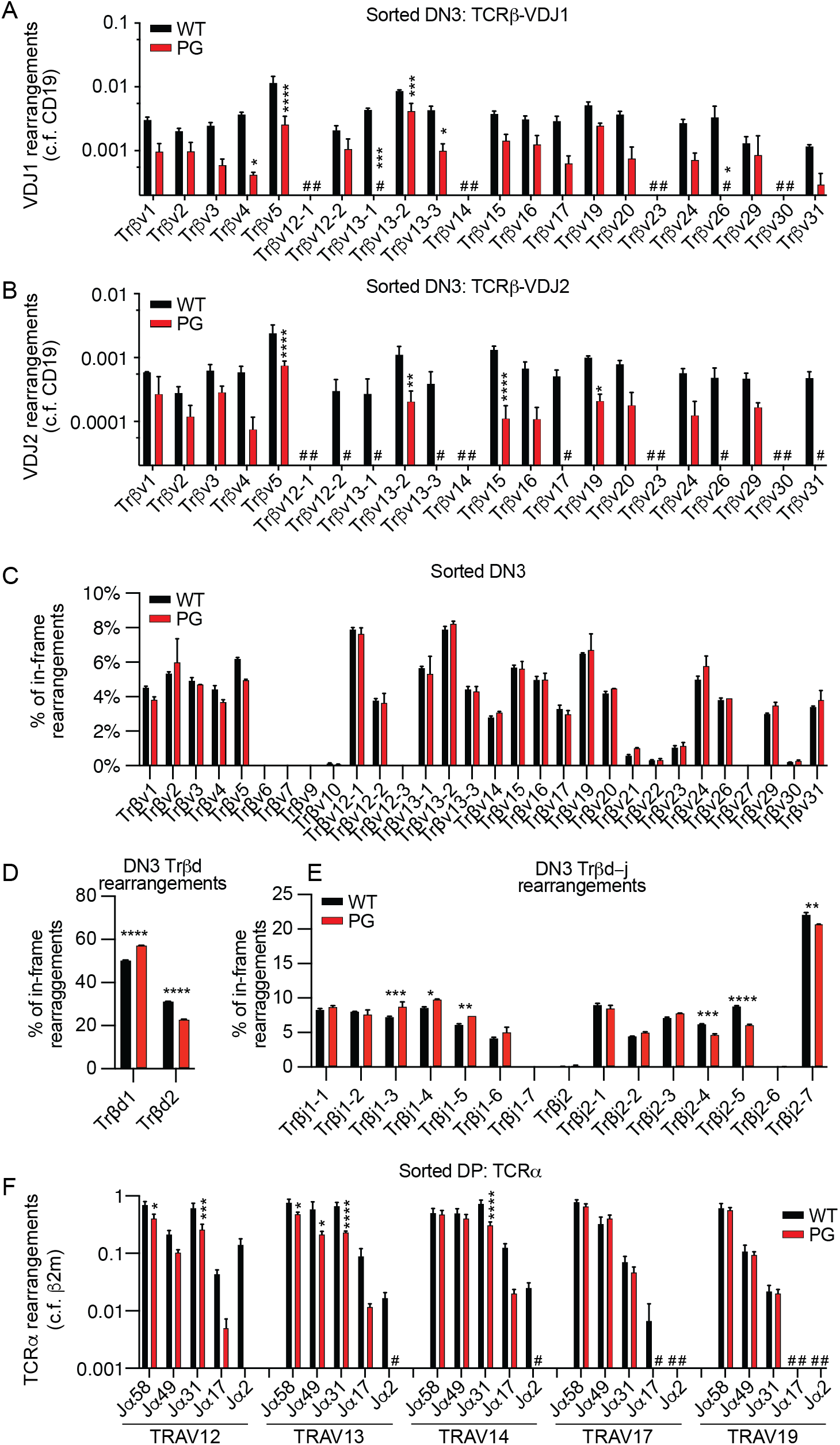
Thymocytes of *PG* mice support diminished levels of *Tcrb* and *Tcra* recombination. **(A-B)** Taqman qPCR quantification of rearrangements of each Vβ segment to DJβ1.1 (A) or DJβ2.1 (B) complexes conducted on genomic DNA from DN3 thymocytes. Signals from each assay were normalized to values from an assay for the invariant CD19 gene. Shown are average values +/- SEM from three independent DN3 thymocyte preparations, each from a different mouse. **(C-E)** Adaptive Immunoseq quantification of the percentage usage of each Vβ (C), Dβ (D), or Jβ (E) segment in *Tcrb* rearrangements of DN3 thymocytes. Shown are average values from two independent DN cell preparations. **(F)** qPCR quantification of specific Vα/Jα rearrangements peformed on genomic DNA from DP thymocytes. Signals from each assay were normalized to values from an assay for the invariant β2m gene. Shown are the average values +/- SEM from three independent DP cell preparations. For all graphs in figure: #, not detected, n.s. p>0.05; * p<0.05, **p<0.01, *** p<0.001, **** p<0.0001

Notably, the degrees to which TCR gene rearrangements and αβ T cell development are reduced in *PG* mice are dramatically less severe than reported for homozygous Rag1 C235Y mutation (14). For example, Rag1^P236G^ mice generate 90% of the normal number of DP thymocytes, whereas homozygous Rag1^C235Y^ mice produce only 1% of the normal number of DP thymocytes. The markedly different severities of these phenotypes suggest that abrogation of Rag1 ubiquitin ligase activity without substantial disruption of Rag1 protein structure only slightly impairs V(D)J recombination and αβ T cell development. Notably, the Rag1^P326G^ protein is expressed at modestly higher levels in thymocytes than wild-type Rag1 protein (19), indicating that reduced Rag1 protein expression levels do not contribute to the slightly impaired development of αβ T cells in *PG* mice. As we were not able to detect Rag1 ubiquitin ligase activity in *WT* mice, we could not confirm the expected abrogation of Rag1 ubiqutin ligase activity in *PG* mice that was observed with Rag1^P236G^ mutation *in vitro*, leaving open the possibility that the mutant Rag1^P326G^ protein exhibits some level of ubiquitin ligase activity *in vivo*.

To confirm that the impaired development of αβ T cells in PG mice is due to cell intrinsic properties, we made and analyzed competitive bone marrow chimeric mice. These data show no signficiant differences in contributions of *WT* cells and *PG* cells at each of DN thymocyte stages (Supplemental Figure 1A). In marked contrast, there were greater contributions of *WT* cells at the DP and both SP thymocyte stages and within the CD4^+^ and CD8^+^ splenic αβ T cell populations (Supplemental Figure 1A, B). The data support the notion that the decreased progression of Rag1^P326G^ thymocytes beyond the DN stage is due to cell intrinsic alterations, most likely from the decreased assembly of TCRβ genes.

### Reduced *Tcrb* gene assembly impairs DN thymocyte development in homozygous Rag1 P326G mice

The greater than normal number of DN3 thymocytes in *PG* mice is not typical of a mutation that impairs V(D)J recombination and could reflect altered β-selection. Thus, to distinguish potential contributions of reduced *Tcrb* gene assembly versus decreased pre-TCR-signaling, we conducted a more refined analysis of DN stage cells based on CD28 protein expression. After successful *Tcrb* gene assembly in CD28^−^CD25^high^ DN3a thymocytes, pre-TCR signaling drives differentiation of DN3b and then DN3c thymocytes characterized by upregulation of CD28 and downregulation of CD25 (22). Mirroring relative levels of *Tcrb* rearrangements in DN3 thymocytes, we observed decreased progression of DN3a cells to the DN3b and DN3c stages, as well as a smaller fraction of DN3a cells expression TCRβ protein, in *PG* mice relative to *WT* mice (Fig. 3, A - F). We compared the percentage of DN3a cells expressing TCRβ protein to the percentage of these cells that develop into DN3b cells. We found equivalent values between *PG* and *WT* mice (Fig. 3, G). We also compared the percentage of total DN3 cells expressing TCRβ protein to the fraction of these cells that progress to the DN4 stage. Here again we found equivalent values between *PG* and *WT* mice (Supplemental Figure 2). Together, these data provide no evidence for diminished pre- TCR signaled development of β-selected DN3 thymocytes. Therefore, the most plausible explanation for impaired DN thymocyte development in *PG* mice is decreased levels of TCRβ gene rearrangements.

**Figure 3.**
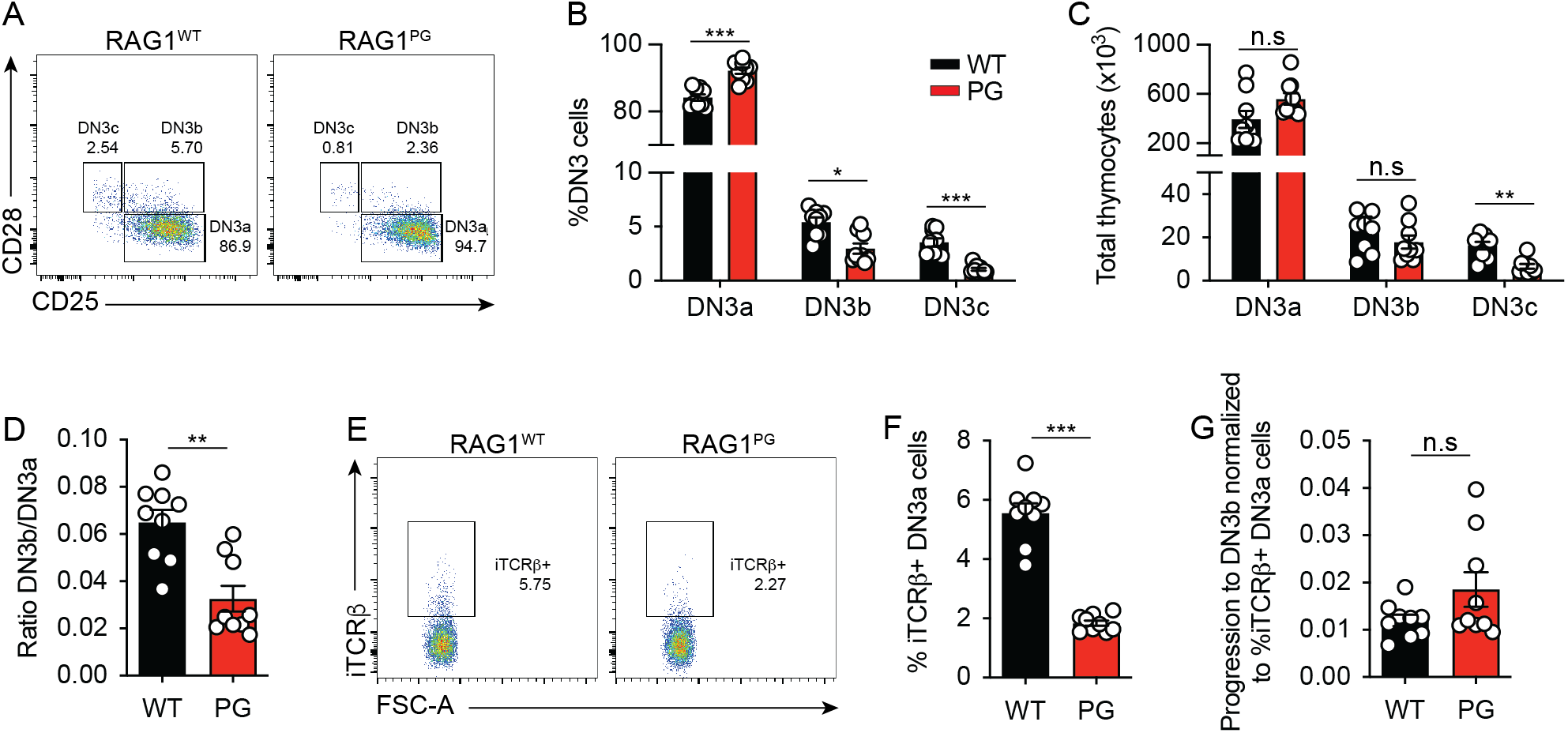
*PG* DN3 thymocytes exhibit normal efficiency of pre-TCR signaled differentiation. **(A)** Representative flow cytometry plots of DN3a, DN3b, and DN3c stages of thymocyte development showing the frequency of cells at each stage. **(B-C)** Quantification of percentages (E) or numbers (F) of total DN3 cells in each stage. Shown are all data points and average values +/- SEM from three independent experiments each with three mice of each genotype (n = 9 mice per genotype, analyzed by multiple t tests using the Holm-Sidak method; n.s. p>0.05, * p<0.05, ** p<0.005, **** p<0.00005). **(D)** Quantification of percentages of DN3a cells that progress into DN3b cells. Shown are all data points and average values +/- SEM from three independent experiments each with three mice of each genotype (n = 9 mice per genotype, analyzed by unpaired t-test with Welch’s correction; **p<0.005). **(E)** Representative flow cytometry plots of intracellular TCRβ protein expression in DN3a thymocytes. **(F)** Quantification of percentages of DN3a cells expressing intracellular TCRβ protein. **(G)** Ratios comparing the percentages of DN3a cells expressing intracellular TCRβ protein to the percentages of DN3a cells that transition into DN3b cells. (**E-J**) Shown are all data points and average values +/- SEM from three independent experiments each with three mice of each genotype (n = 9 mice per genotype, analyzed by unpaired t-test with Welch’s correction; n.s. p>0.05, *** p<0.0005).

### Homozygous Rag1 P326G mutation decreases the efficiencies of positive and negative selection of conventional αβ T cells and differentiation of iNKT cells

In developing B cells, RAG DSBs generated during *Igk* gene assembly signal transcriptional regulation of proteins that govern antigen receptor signaling, lymphocyte selection, cellular survival, and/or cellular proliferation (23–25). Considering this and the decreased *Tcra* gene assembly in *PG* DP thymocytes, we investigated whether any of the thymocyte selection and subsequent differentiation processes that depend on αβ TCR signaling might be impaired in *PG* mice. As a proxy to evaluate positive selection in *PG* mice, we determined the frequency of αβ TCR expressing (TCRβ^+^) Stage 1 thymocytes that initiate positive selection as indicated by progression to Stage 2 in *PG* and *WT* mice (Fig. 4 A). By analyzing only αβ TCR expressing cells, we circumvent potential effects of decreased *Tcra* gene rearrangements reducing the fraction of DP thymocytes that express αβ TCRs and are subject to selection. This frequency is lower in *PG* mice (Fig. 4 B), consistent with modestly less efficient positive selection of αβ TCR expressing DP thymocytes upon homozygous Rag1^P326G^ mutation. Yet, we note that this is an imperfect means to quantify positive selection because it cannot account for possible differences in progression through subsequent stages or cellular fate decisions from specific TCR/pMHC interactions. Thus, we also quantified numbers of cells at each of the four stages of positive selection, finding fewer cells at each stage in *PG* mice relative to *WT* mice (Fig. 4 C). Importantly, there is a greater decrease of cellularity in stage 2 (1.71x) as compared to stage 1 (1.25x), indicating fewer cells are moving from stage 1 to stage 2 in *PG* mice. Finally, we quantified CD69 expression and the ratio of CD69 to TCRβ expression at each stage as CD69 upregulation correlates directly with αβ TCR signaling strength. Although we observed no difference of absolute levels of CD69 expression, *PG* mice have decreased CD69 expression relative to TCRβ expression in stage 2 thymocytes that are initiating positively selection (Fig. 4 D). Collectively, these analyses indicate that αβ TCR signaled upregulation of CD69 and positive selection is less efficient in *PG* mice. However, they do not distinguish between potential effects from altered primary repertoires of *Tcrb* and *Tcra* genes versus diminished intrinsic αβ TCR signaling capacity. They also do not address the possibility that the <10% reduction in numbers of total thymocytes in *PG* mice alters thymic architecture in a manner that decreases the fraction of DP cells that engage pMHC presented by TECs or DCs.

**Figure 4.**
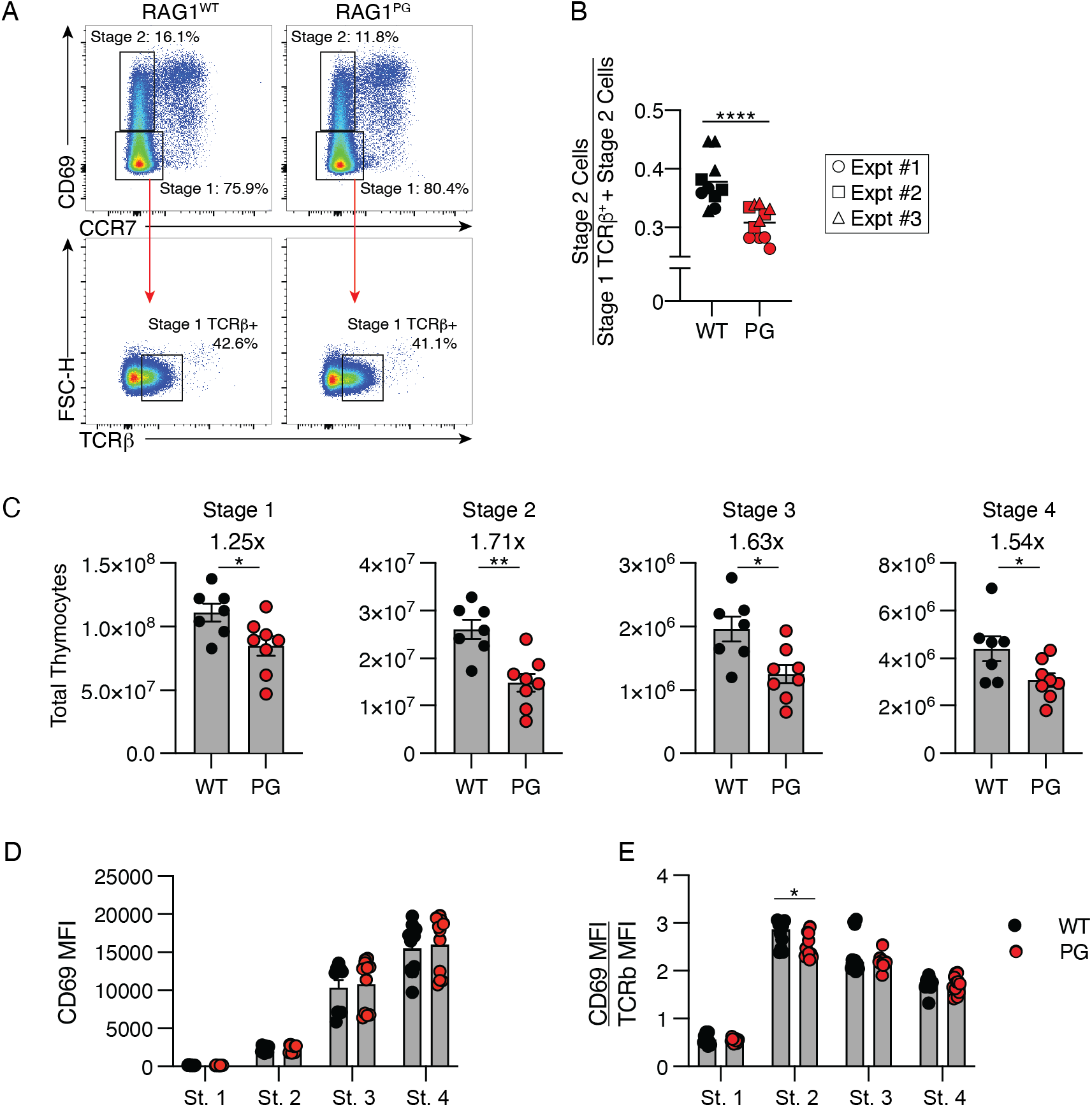
*PG* mice exhibit impaired positive selection. **(A)** Gating strategy for measuring positive selection of thymocytes showing Stage 1 and 2 gates. The TCRβ^+^ gate was set using wild-type thymocytes by gating on live, singlet, CD4^+^CD8^+^ cells, preparing a histogram showing TCRβ staining on cells within this gate, and gating on the shoulder that represents cells expressing TCRβ. **(B)** Quantification of the ratio of the number of Stage 2 cells to the total number of Stage 1 and 2 cells. Shown are all data points and global mean from three independent experiments (n = 10-11 mice per genotype, 2-way ANOVA indicating effect of genotype). **(C)** Quantification of numbers of cells in each stage of positive selection. Dots indicate individual mice. Data pooled from 2-independent experiments (n = 7-8 mice per genotype, bars indicate mean +/- SEM analyzed by Man-Whitney test). (**D**) Quantification of CD69 expression alone and in comparison to TCRβ expression on cells at each stage of positive selection. Dots indicate individual mice. Data pooled from 2-independent experiments (n = 8 mice per genotype, bars indicate mean +/- SEM. Analysed by 2-way ANOVA and Sidak multiple comparison post-test) For all graphs in figure: n.s. p>0.05; * p<0.05, **p<0.01, *** p<0.001, **** p<0.0001.

To determine whether the Rag1^P326G^ mutation impairs negative selection , we utilized a naturally occurring system of superantigen-mediated negative selection (26). Laboratory mice express remnant mouse mammary tumor virus (MMTV) retroviral products. Some of these products can bind the MHCII I-E^d^ protein and certain Vβ peptides, including Vβ5.1-5.2 (encoded by *Trbv12.1*-*12.2*) and Vβ12 (encoded by *Trbv15*), to trigger exceptionally strong αβ TCR signaling that promotes negative selection (26, 27). Our sequencing analysis of *Tcrb* gene rearrangements in DN3 thymocytes shows equivalent representation of *Trbv12.1*, *Trbv12.2*, and *Trbv15* in the primary Vβ repertoire (Fig. 2 C). Considering that Vβ repertoire remains constant during DN-to-DP thymocyte development (28), assaying superantigen-mediated deletion of Vβ5.1-5.2 or Vβ12 accounts for differences of Jβ repertoire of *Tcrb* genes assembled in *WT* and *PG* mice. Moreover, the reduced level and altered repertoire of *Tcra* genes assembled in DP cells of *PG* mice would not influence superantigen-mediated deletion of thymocytes expressing αβ TCRs with specific Vβ peptides. BALB/c, but not C57BL/6, strain mice express I-E^d^ protein. As a result, thymocytes expressing Vβ5.1-5.2 or Vβ12 are efficiently deleted in BALB/c, but not B6, genetic backgrounds. We bred BALB/c background *Rag1^-/-^* mice with C57BL/6 background Rag1 *PG* or *WT* mice to generate and analyze *Rag1^PG/-^* and *Rag1^WT/-^* mice expressing I-E^d^ protein. To ascertain whether these mice exhibit a difference in superantigen-mediated negative selection, we measured the frequencies of DP thymocytes that have not completed αβ TCR selection and post-selection SP thymocytes expressing I-E^d^:MMTV- reactive (Vβ5^+^ or Vβ12^+^) or -unreactive (Vβ6^+^, Vβ8^+^, or Vβ14^+^ encoded by *Trbv19*, *Trbv13.1-13.3*, or *Trbv31*, respectively) αβ TCRs (Fig. 5, A-F). We calculated the percentages of DP and SP thymocytes expressing each Vβ peptide (Fig. 5, B-F). Our analysis indicates that greater fractions of I-E^d^:MMTV- reactive thymocytes proceed through DP-to-SP development in *Rag1^PG/-^* mice as compared to *Rag1^WT/-^* mice (Fig. 5, B, C, and G), consistent with impaired negative selection of highly self-reactive thymocytes. In contrast, we found that the types of un-reactive thymocytes assayed develop equivalently in each genotype (Fig. 5, D -G). While the latter finding does not reflect less efficient positive selection, we note that this experiment monitors a small subset of thymocytes expressing specific Vβ peptides and thus is not as sensitive as determining the frequencies of overall DP thymocytes that initiate positive selection. Regardless, the data indicate that *PG* mice exhibit less efficient deletion of thymocytes that express αβ TCRs containing superantigen-reactive Vβ peptides. As Vβ repertoire is normal in *PG* mice (Fig. 2C), plausible explanations for this impaired form of negative selection include that intrinsic αβ TCR signaling capacity is dampened and/or reduced numbers of DP thymocytes prevents some from contacting or maintaining interaction with superantigen.

**Figure 5.**
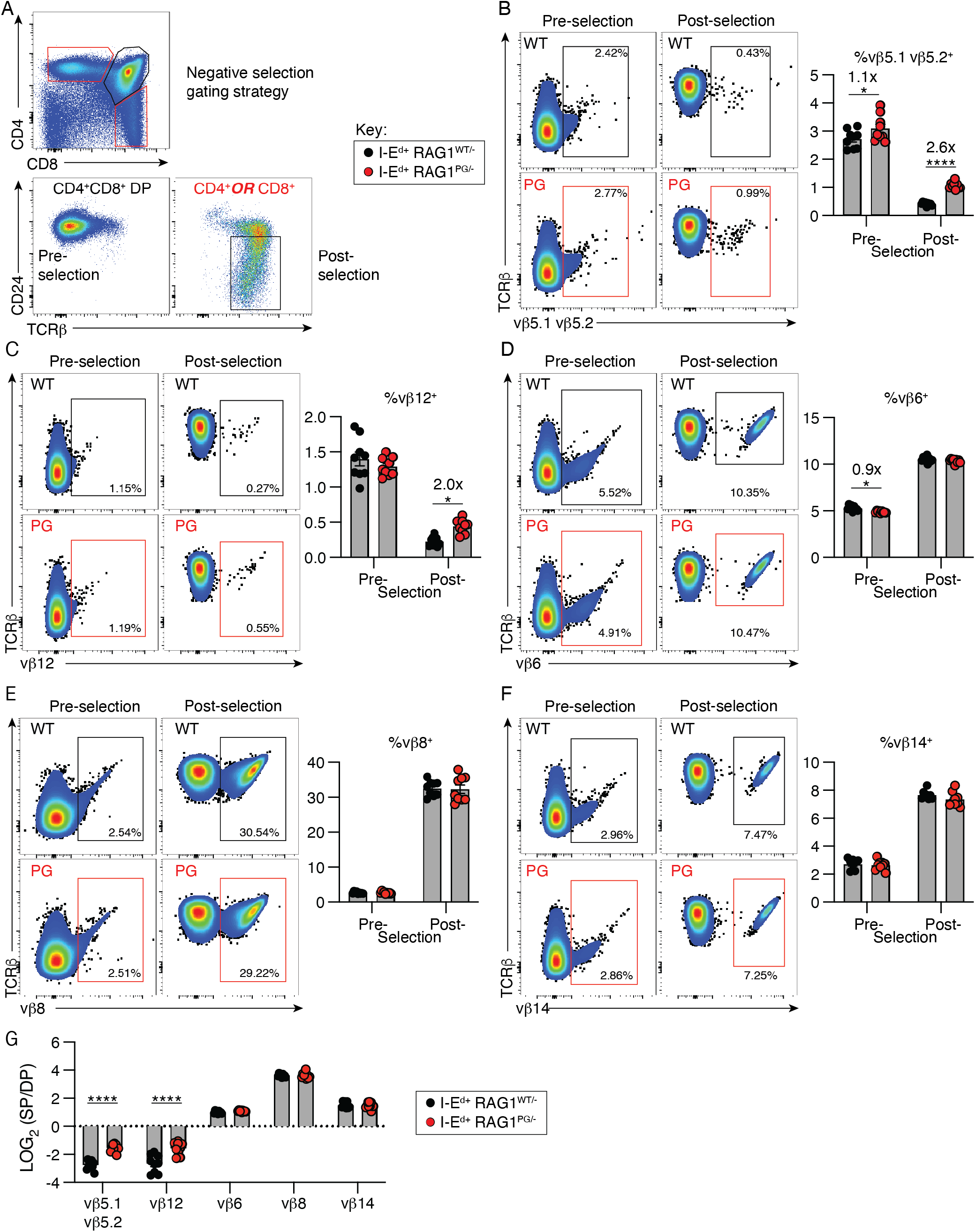
*PG* mice exhibit impaired superantigen-mediated negative selection. **(A)** Gating strategy form measuring negative selection of specific Vβ^+^ thymocytes in F1 progeny of C57BL/6 background *WT* or *PG* mice crossed with BALB/c background *Rag1^-/-^* mice. Pre-gated on live singlets. **(B-F)** Representative analyses and quantification of the indicated Vβ^+^ cells in DP thymocytes that have not completed αβ TCR selection or post-selection SP thymocytes. **(G)** Data from B-F displayed as LOG_2_ ratio of respective percent Vβ^+^ cells in SP versus DP gates. Shown are all data points and average +/- SEM from two independent experiments (n = 9 mice per genotype 2-way ANOVA with Sidak multiple comparison post- test). For any significantly different combinations, fold change is indicated. For all graphs in figure: * p<0.05, **** p<0.0001.

Finally, to ascertain potential effects of Rag1^P326G^ mutation on agonist selection, we focused on assaying iNKT cell development because their limited TCR diversity and specificity facilitates their identification and assays of their selection and differentiation subsequent to V(D)J recombination. This lineage develops when TCRs containing an invariant Vα14-Jα18 TCRα chain and TCRβ chains with Vβ regions encoded by *Trbv1*, *Trbv13.1-13.3*, or *Trbv19* bind lipids presented by the MHC-like CD1d protein on different DP thymocytes (29–31). These interactions induce strong TCR signals that mediate agonist selection of DP cells into αβ TCR-expressing iNKT common progenitor (Stage 0) cells and their differentiation into iNKT precursor (Stage 1) cells and then into different mature effector subsets (iNKT1, iNKT2, or iNKT17)(32). We first used a CD1d tetramer loaded with PBS57, an analogue of the α-galactosylceramide (α-GalCer) glycolipid, to identify α-GalCer-reactive iNKT lineage cells in *WT* and *PG* mice. Unloaded “empty” tetramer showed little to no staining in any of the samples (exemplified in Fig. 6A). As compared to *WT* mice, *PG* mice had fewer absolute numbers of thymic α-GalCer-reactive iNKT lineage cells (Fig. 6, A and B). While *PG* mice also tended to have fewer splenic iNKT cells (Fig. 6, A and B), this difference did not reach statistical significance (p=0.06) with the numbers of mice analyzed. We used Taqman PCR to show that the levels of Vα14-to-Jα18 rearrangements in sorted DP thymocytes of *PG* mice are ∼10% relative to *WT* mice (Fig. 6 C), which is consistent with reduced generation of Vα14-Jα18 TCRα chains contributing to the lower numbers of thymic α-GalCer-reactive iNKT cells in *PG* mice. However, we observed no significant differences in the numbers of PBS57 loaded tetramer-staining cells at both Stage 0 (CD24^hi^CD44^−^NK1.1^−^) or Stage 1 (CD24^lo^CD44^−^NK1.1^−^) in *WT* and *PG* mice (Fig. 6, D and E), indicating that the reduction of Vα14-to-Jα18 rearrangements in *PG* DP thymocytes does not translate into decreased agonist selection of α-GalCer-reactive iNKT lineage cells. In contrast, we observed that *PG* mice have lower numbers of NKT2/17 (CD24^lo^CD44^+^NK1.1^−^) and NKT1 (CD24^lo^CD44^+^NK1.1^+^) effector iNKT cells that develop from Stage 1 precursor cells (Fig. 6, D and E). Notably, the median fluorescence intensities of TCRβ expression and PBS57/CD1d tetramer binding of thymocytes was similar or slightly higher in *PG* mice relative to *WT* mice (Fig. 6 A, data not shown), indicating that α- GalCer-reactive iNKT cells of *PG* mice have a normal range of intrinsic affinity for this lipid presented by CD1d. We also analyzed iNKT populations in the competitive setting of mixed bone marrow chimeras comparing mice that received donor bone marrow that was either 100% *PG* or at 1:1 mix of *PG* and *WT* marrow. The *WT* cells in the mixed chimeras outcompeted the *PG* cells starting at Stage 1 with more pronounced effects in the terminally differentiated effector iNKT cell types (Supplemental Figure 1). The same effect can be seen from the few remnant *WT* cells in the 100% *PG* recipient mice (Supplemental Figure 1, C). These data indicate that there is a cell-intrinsic loss of fitness in iNKT cell development in *PG* cells as they progress from Stage 0 to Stage 1 that is not revealed in the non-competitive environment. Because the fitness of the *PG* iNKT Stage 0 and 1 cells is poor in the 1:1 mixed environment, it is difficult to assess cell-intrinsic iNKT differentiation in this setting. Yet, the few remnant *WT* cells in the 100% *PG* recipient mice continue to incrementally outcompete *PG* cells in both NKT1 and NKT 2/17 differentiation, demonstrating a cell-intrinsic differentiation defect in this system. The basis for this defect could include altered αβ TCR signaling arising from differences in Dβ-, Jβ-, CDR3β-, or CDR3α-encoded amino acids, diminished RAG DSB-signalled changes in expression of proteins that control iNTK cell differentiation, or both.

**Figure 6.**
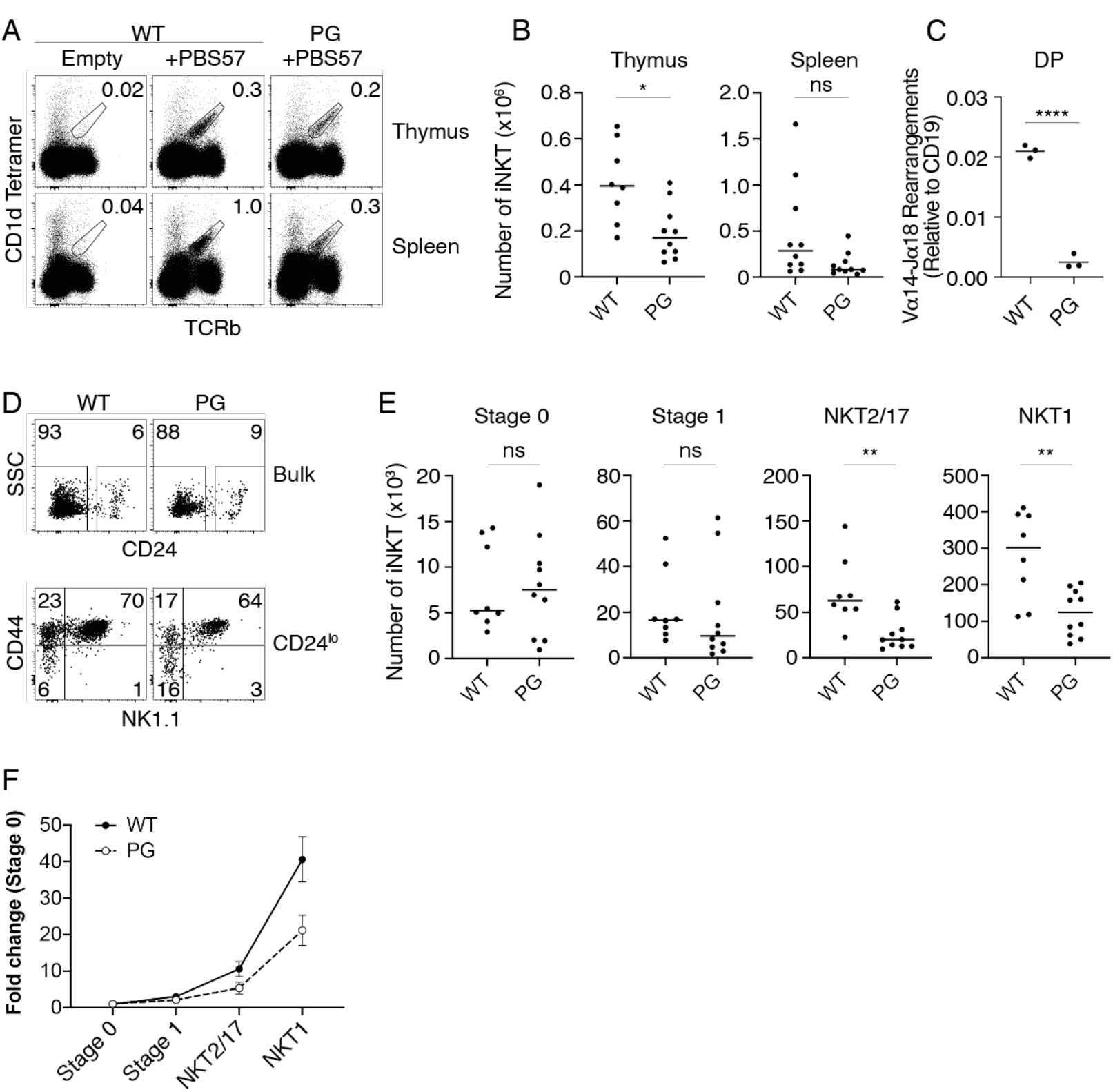
*PG* mice have impaired agonist selection of iNKT cells. (**A-B)** Representative flow cytometry plots (**A**) and quantification (**B**) of PBS57/CD1d tetramer binding iNKT cells in thymuses or spleeens. Gated on live, singlet lymphocytes. “Empty” denotes unloaded tetramer staining. (**B**) Data from three independent experiments with 1-4 mice per group. Mann-Whitney test: *, p=0.01 (thymus); ns, not significant (spleen). (**C**) Taqman PCR quantification of Vα14-Jα18 rearrangements from sorted DP thymocytes. The data were generated from three technical replicates of three independent biological samples from mice of each genotype. Data are represented as Mean +/- SEM analyzed by an unpaired parametric t test with Welchs correction. p<0.001. (**D-E)** Representative flow cytometry plots and gating strategy (**D**) or quantification (**E**) of Stage 0 (CD24^hi^ CD44-NK1.1-), Stage 1 (CD44-NK1.1-), NKT2/17 (CD44+NK1.1-), and NKT1 (CD44+NK1.1+) iNKT cells.

### Mature αβ T cells of homozygous Rag1 P326G mice exhibit normal responses to activation

RAG endonuclease activity during lymphocyte ontogeny triggers permanent changes in gene expression that correlate with impaired responses of mature lymphocytes (33). Considering that RAG endonuclease function is diminished in thymocytes of *PG* mice, we sought to elucidate whether the mature αβ T cells of these animals have normal or altered response to activation. We first conducted *in vitro* analyses. Upon stimulation with PMA and ionomycin, splenic CD44^hi^ αβ T cells from *PG* mice produce normal levels of IFNγ, TNFα, IL-2, and IL-17A (Fig. 7 A). These cells also show normal proliferation upon anti-CD3/anti- CD28 stimulation (Fig. 7 B). Moreover, stimulated CD4^+^ and CD8^+^ αβ T cells of *PG* mice normally express their canonical transcription factors (Fig. 7, C and D) and cytokines (Fig. 7, E and F) when cultured in respective skewing conditions. We next monitored αβ T cell responses *in vivo* following infection of mice with an acute strain of lymphocytic choriomeningitis virus (LCMV-Armstrong). We analyzed mice either seven days later to monitor at the immune response peak or 35 days later to monitor memory cells. At each timepoint, we detected similar frequencies and numbers of antigen-specific CD8^+^ αβ T cells that stain with gp33:H-2Db tetramer in *PG* and *WT* mice (Fig. 7 G), indicating that *PG* mice display normal numbers and dynamic responses of αβ T cells that recognize LCMV-Armstrong. We also observed equivalent dynamic expression of Tbet, Eomes, KLRG1, and CD127 proteins on αβ T cells from *PG* and *WT* mice (Fig. 7 H), indicating normal contraction of short-lived effector cells (SLECs), and sustained memory/memory precursor effector cells (MPECs) in *PG* mice. Finally, we detected similar numbers of antigen specific αβ T cells able to respond to the gp33, NP396, or gp61 viral peptides within a peptide re- stimulation assay (Fig. 7, I and J). Together, these data demonstrate that the mature αβ T cells of *PG* mice have normal responses to activation, in the tested conditions.

**Figure 7.**
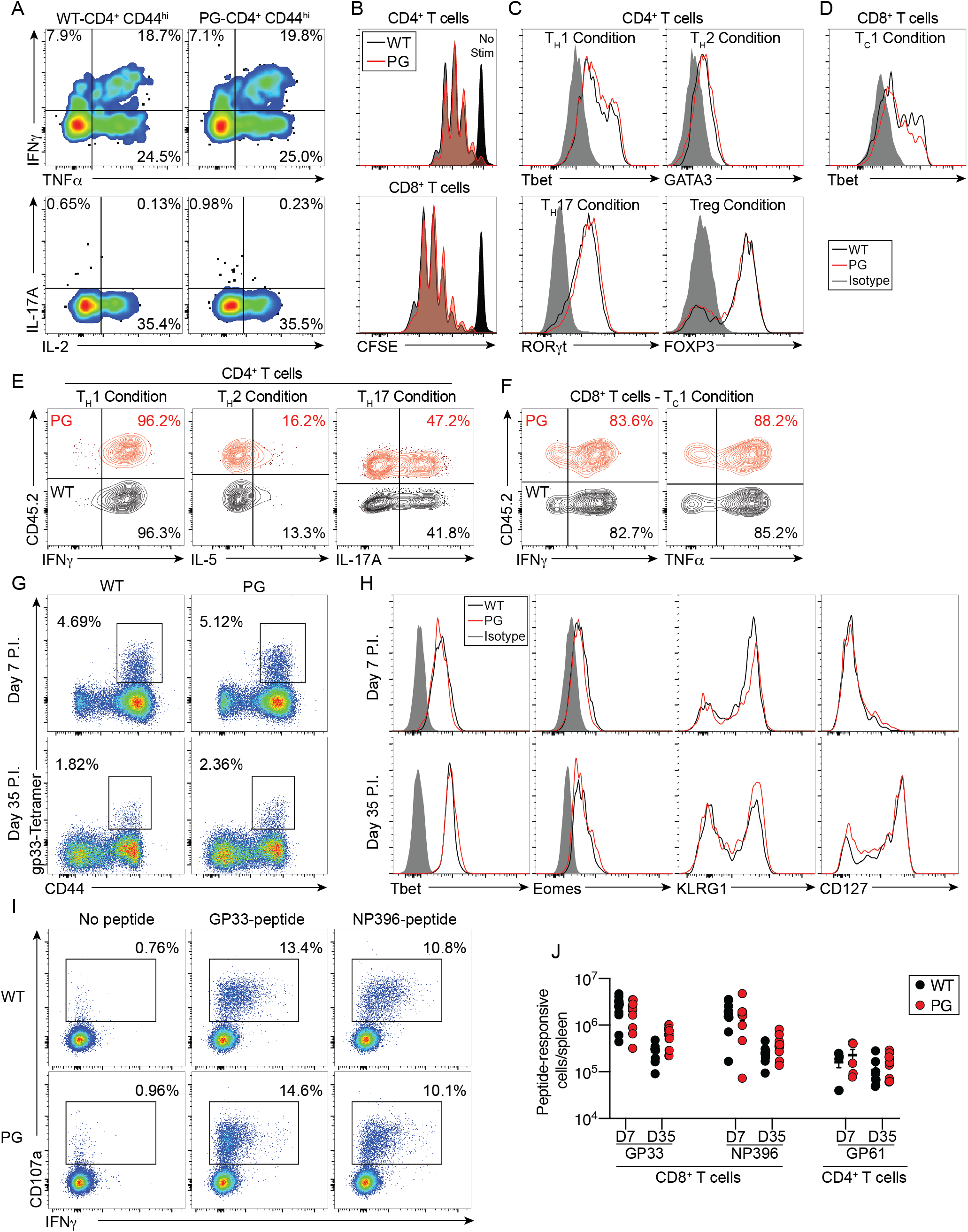
Mature αβ T cells of PG mice exhibit normal immune response. **(A)** Representative plot of cytokine expression on CD4^+^CD44^hi^ αβ T cells following *ex vivo* stimulation of splenocytes with PMA and ionomycin. **(B-F)** Representative analyses of naïve splenic CD4^+^ or CD8^+^ T cells after *ex vivo* simulation with anti-CD3/anti-CD28 antibodies and additional antibodies to skew towards indicated lineages. Shown is CFSE incorporation (B) or expression of relevant transcription factors (C, D) or cytokines (E, F). **(G, H)** Representative flow data showing (G) frequencies of splenic αβ T cells that stain with the gp33:H-2Db tetramer or (H) express indicated cytokines after LCMV infection of mice. Data are representative of two independent experiments with at least four mice of each genotype. **(I, J)** Representative plots (J) and (I) quantification of antigen-specific αβ T cells able to respond to indicated viral peptides within a peptide re-stimulation assay. (A-F) Data are representative of two independent experiments. (G-J) Data are combined from four independent experiments (two D7 and two D35 timepoints) with 8-11 mice of each genotype per time-point.

### Mature αβ T cells of homozygous Rag1^P326G^ mice exhibit increased autoimmune potential

Within a setting of less efficient negative selection as with homozygous Rag1^P326G^ mutation, an increased predisposition to αβ T cell mediated autoimmunity would be expected. However, we did not observe overt autoimmune symptoms within *PG* mice, possibly because mice of the C57BL/6 background are resistant to spontaneous autoimmune diseases (34, 35). Thus, to investigate whether the Rag1^P326G^ mutation can predispose to autoimmunity, we utilized an induced model of autoimmune colitis in C57BL/6 mice.

The colitis model involves transferring into *Rag1^-/-^* mice naïve CD4^+^ effector αβ T cells, which become activated and trigger cytokine-driven colitis and associated systemic pathologies including weight loss due to the absence of Tregs (36–38). We observed accelerated wasting of *Rag1^-/-^* mice receiving *PG* cells relative to *WT* cells or no transfer (Fig. 8 A). We analyzed mice at eight weeks after transfer. As compared to *WT* recipients, we detected greater numbers of total splenic CD4^+^ T cells and of CD4^+^ T cells producing IFNγ, TNFα, and/or GM-CSF cytokines (Fig. 8, B and C). Colon length is used as a measure of disease severity in this model, and mice receiving transferred *PG* cells had shorter colons relative to mice receiving *WT* cells, consistent with worse disease caused by *PG* cells (Fig. 8 D). Histologic assessment and scoring in this colitis model has been formalized (38), and can be grossly broken down into scores related to inflammatory cell infiltration and scores related to injury of tissue architecture. There was no statistical difference in inflammation scores between mice receiving *WT* and *PG* cells. However, tissue damage scores were higher in mice receiving *PG* cells, suggesting that, on a per cell basis, *PG* cells induce greater tissue injury in this model (Fig. 8, E-F). Notably, specifically in *PG*-transferred animals, we observed autoimmune pathologies in addition to colitis, including dermatitis, chylous ascites, severe small intestine enlargement, and neurological distress (Fig. 8 H). Dermatitis was most common and involved massive inflammatory infiltration and tissue damage (Fig. 8 I). Consistent with more widespread autoimmune conditions in *PG*-transferred mice, we observed significant splenomegaly in these mice (Fig. 8 J). Finally, while not statistically significant, we observed an increased rate of premature death in the mice transferred with *PG* cells compared to *WT* cells (Fig. 8 K), which was especially surprising because death is an unexpected outcome in this model. Collectively, the data from this colitis model indicate that CD4^+^ effector T cells from *PG* mice have greater than normal intrinsic autoimmune potential.

**Figure 8.**
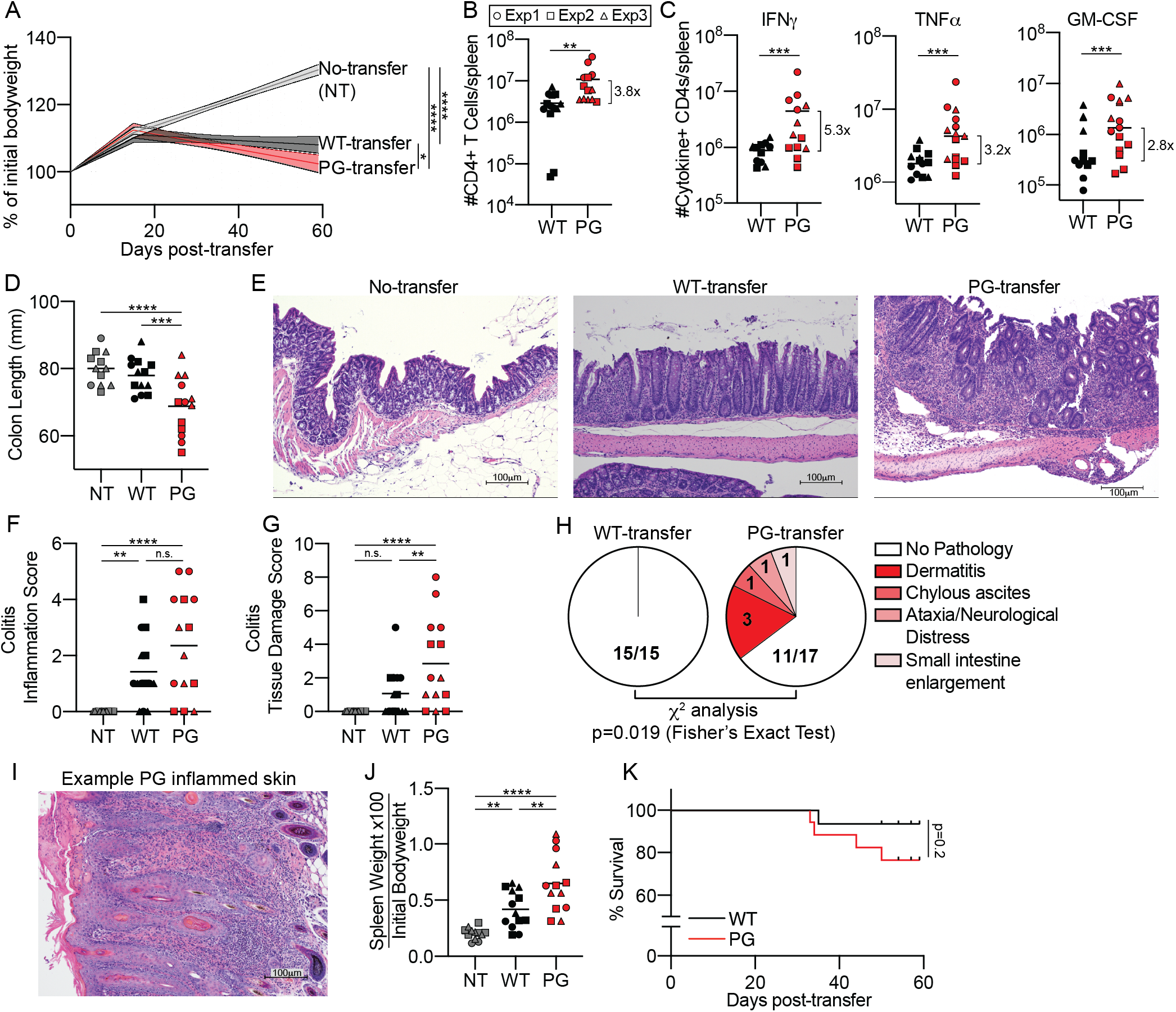
Mature αβ T cells of *PG* mice possess higher than normal autoimmune hazard**. (A)** Graph of body weights of *Rag1^-/-^* hosts over time after transfer of *WT* or *PG* CD4^+^ T cells or no cells (NT). Data combined from three independent experiments with a total of 13-17 mice of each genotype. Lines indicate fitted segmented-linear regression lines +/- 95% confidence interval. Day 15 was fixed as the inflection point. Difference between gradients of second slope was determined by linear regression analysis. **(B)** Numbers of CD4^+^ T cells in spleen eight weeks post-transfer. **(C)** Total number of IFNγ, TNFα, and/or GM-CSF producing cells per spleen. Shown are all data points from three independent experiments with 12-14 mice per group. Bar indicates global mean. Pre-gating on live singlets, CD90.2^+^, CD4^+^, and CD44^+^. **(D)** Colon length of mice sacrificed at eight weeks post-infection. **(E)** Representative hemotoxylin and eosin stained colon sections from respective mice at eitght weeks post-transfer. 10x magnification. **(F-G)** Total colitis score as assessed by blinded pathologist. Scored as per previously described guidelines for the five phenotypes (38): (F) Colon inflammation score (contribution of scores for crypt abcesses and inflammatory cell infiltrate). (G) Colon tissue damage score (contribition of scores for crypt architecture, tissue damage, goblet cell loss). **(H)** Non-colitis pathologies described for mice transferred with *WT* versus *PG* cells. **(I)** Hemotoxylin and eosin stained skin section from *PG*-transferred mouse with severe dermatitis at eight weeks post-transfer. 10x magnification. **(J)** Spleen weight at eight weeks post-transfer normalized to total mouse bodyweight prior to transfer. **(K)** Survival curves of mice transferred with *WT* or *PG* cells. Data combined from three independent experiments. Statistical analysis = Log-rank Mantel- Cox test. Overall: 0/13 NT mice died, 1/15 *WT*-transferred mice died, and 4/17 *PG*-transferred mice died. (B, C, F, G, and J) Shown are all data points from three independent experiments, 12-14 mice per group. Experiment indicated by shape of data point. Bar indicates global mean. Analyzed by 2-way ANOVA with Tukey HSD post-test. n.s. p>0.05; * p<0.05, **p<0.01, *** p<0.001, **** p<0.0001

## Discussion

Our analysis of *PG* mice demonstrates that the RAG1 RING domain stimulates V(D)J recombination of endogenous TCR loci and resulting αβ T cell development *in vivo*. The decreased V(D)J recombination activity in PG mice manifests as lower levels of Vβ-to-DJβ and Vα-to-Jα rearrangemetns within DN and DP thymocytes, respectively. These reduced Vβ rearrangements diminish overall progression of the DN3 thymocyte population to the DN4 and then DP stage by lowering the frequency of DN3 cells that create TCRβ protein. Similarly, reduced Vα rearrangements likely contribute to less efficient overall positive selection of the DP thymocye population by lowering the frequency of DP cells that establish αβ TCRs. Moreover, the 10% lower numbers of DP thymocytes in *PG* mice resulting from decreased expansion of the DN population could impair positive selection by reducing engagement of αβ TCRs on DP cells with pMHC on TECs or DCs. Our data align with the recent report that Rag1^P326G^ mutation in mice diminishes V(D)J recombination of IgH and Igκ loci and the B cell developmental transitions that depend on these rearrangements by reducing RAG cleavage (19). The Rag1^P326G^ mutation might intrinsically decrease RAG endonuclease activity by altering Rag1 protein structure and/or impairing Rag1-mediated autoubiquitylation of RAG complexes (10, 11, 15). As the RAG1 RING domain promotes ubiquitylation of histone H3 proteins (12–14), our data raise the possibility that the RAG1 ubiquitin ligase might stimulate accessibility of chromatin over TCR and Ig loci to enhance RAG binding, cleavage, or both. Moreover, because RAG1 RING domain mutations that impair histone H3 ubiquitylation result in accumulation of coding ends and signal ends in episomal substrates (13), less efficient V(D)J coding join formation might diminish TCR and Ig gene rearrangements and resulting lymphocyte differentation in *PG* mice. Eludicating the precise molecular mechanisms by which the RAG1 RING domain promotes V(D)J recombination *in vivo* requires biochemical studies of RAG complexes in *WT* and *PG* cells, epigenetic analyses of TCR and Ig loci in mice that express endonuclease-inactivated RAG complexes with or without Rag1^P326G^ mutation, and characterization of the ubiquitylated proteome in all four cell types.

Our analysis of *PG* mice also reveals that the RAG1 RING domain shapes primary TCR gene repertoires and might thereby regulate positive selection and negative selection of conventional αβ T cells. The amino acids encoded by rearranged gene segments or *de novo* CDR3 nucleotoides influence antigen recognition, signaling, and selection of αβ TCRs. Although we did not analyze CDR3 nucleotoides, our data indicate differences in utilization of Dβ and Jβ segments within Vβ-to-DJβ rearrangements and Vα and Jα segments within Vα-to-Jα rearrangements between *WT* and *PG* mice. The altered utilization of TCRβ and/or TCRα gene segments in V(D)J rearrangements of *PG* mice might create a primary TCR repertoire that overall is less efficient at promoting positive selection. Moreover, the altered usage of TCRβ and/or TCRα gene segments in *PG* mice could produce a TCR repertoire with a greater than normal frequency of highly selfreactive receptors that escapes negative selection. Decreased RAG endonuclease activity from Rag1^P326G^ mutation likely reduces the utilization of Dβ2 and 3’Jαs by lowering the frequency of V recombination to these gene segments after V recombination to Dβ1 or 5’Jαs. One prediction of impaired RAG cleavage activity is elevated usage of gene segments with stronger RAG-targeting recombination signal sequence (RSS) elements (20). This is the case for Vβ usage in mice with profoundly low RAG endonuclease activity caused by a deletion of the RING domain and all more N-terminal residues (21). The Jβ2.5 gene segment has a stronger RSS than Jβ2.2 (39). Considering that Jβ2.5 usage increases and Jβ2.2 usage remains normal in TCRβ gene rearrangements of *PG* mice, Rag1^P326G^ mutation likely alters Jβ utilization by means other than reducing RAG endonuclease activity. The mutation could prevent the RAG1 ubiquitin ligase from promoting accessibility of some Jβ RSSs, destablilize snynaptic complexes based on distinct properties of participating Jβ RSSs, or both. The same approach needed to elucidate how the RAG1 RING domain promotes V(D)J recominaiton *in vivo* would provide mechanistic insights into how it also shapes primary antigen receptor gene repertoires.

Although further experimentation is required to better understand how the RAG1 RING domain regulates V(D)J recombination, any discussion of αβ TCR selection and resultant differentiation of DP thymocytes would be incomplete without consideration of a possibilities that the RAG1 RING domain regulates these processes independent of assembling TCR genes. The impaired superantigen-mediated negative selection of conventional αβ T cells and differentiation of iNKT lineage cells in *PG* mice suggest potential functions of the RAG1 RING domain beyond stimulating and shaping TCR gene rearrangements. The molecular basis for superantigen-mediated negative selection is that these antigens interact simultaneously with MHC proteins on TECs and particular Vβ peptides on DP thymocytes to illicit very strong TCR signaling that triggers apoptosis. To our knowledge, there is no evidence that Dβ-, Jβ-, CDR3β-, Vα, Jα, or CDR3α- encoded amino acids influence superantigen-directed negative selection. Thus, as Vβ usage within Vβ-to- DJβ rearrangements is normal in *PG* mice, the altered Dβ, Jβ, Vα, and Jα usage should not impair superantigen-driven negative selection of DP thymocytes that express relevant Vβ peptides within their αβ TCRs. How iNKT cell subsets develop from agonist-selected iNKT progenitor cells remains incompletely understood. A recent study in mice expressing αβ TCRs from a single pre-assembled Vα14- Jα18 rearrangement formulated a model that iNKT cell development is a two step process where the first step involves transient uniform TCR signaling that drives agonist selection of DP thymocytes into iNKT common precursors and the second step involves antigen/TCR-independent differentiation of different mature effector subset through differential gene expression (40). However, this study did not investigate the effects of TCR signaling strength on differentiation of iNKT precursor cells. Prior work that the development of NKT2 and NKT17 but not NKT1 cells is impaired by a hypomorphic mutation of the Zap70 TCR signaling protein that weakens TCR signaling supports the notion that strength of TCR signaling influences iNKT cell differentiation (40). Although TCRβ amino acids encoded by Jβ segments and CDR3β nucleotides influence αβ TCR specifity of iNKT cells (41), it is hard to explain how the minor changes in Dβ and Jβ usage in *PG* mice would substantially alter TCR signaling to impair the generation of iNKT subtypes from iNKT precursor cells.

We envision two not-mutually exlusive possibilities that account for the impaired superantigen-mediated negative selection of conventional αβ T cells and differentiation of NKT precursor cells in *PG* mice. One is that the 10% reduction in DP thymocyte numbers decreases frequencies or durations of αβ TCR contacts with pMHC on other thymic cells or CD1d/lipid complexes on other DP cells, causing weaker or otherwise different TCR signaling. The second possibility is that Rag1^P326G^ mutation modifies the expression and/or activities of proteins in DP thymocytes and derivative iNKT precursor cells. To explore this latter scenario, we quantified Zap70 expression because reduced Zap70 protein levels and/or activity impairs positive and negative selection (42–44), as well as iNKT cell differentiation (45, 46). We detected lower levels of Zap70 protein but not mRNA in DP cells of *PG* mice at each of the four stages of αβ TCR-signaled positive selection as defined by CD69 and CCR7 protein expression (Supplemental Figure 3). Zap70 proteins levels are normal in splenic αβ T cells of *PG* mice, correlating with a time point when *Rag1* is not expressed and therefore should not be able to directly influence protein levels (Supplemental Figure 3). We did not measure Zap70 expression in iNKT lineage cells. The decreased Zap70 expression in DP thymocytes might contribute to the impaired positive selection and superantigen-mediated negative selection of *PG* mice. It also could impact generation of the NKT2/17 cell population. Furthermore, it could contribute to relative lack of fitness in of *PG* iNKT cells progressing from Stage 0 to Stage 1 when competing with wild type cells. Independent of whether these Zap70 differences are causative of the observed altered negative selection and NKT differentiation in *PG* mice, reduced Zap70 expression demonstrates proof-of-concept that the Rag1^P326G^ mutation alters protein expression in developing αβ T cells as suggested in our second hypothesis above. RAG DSBs induced in pre-B cells signal temporal changes in gene transcription to modulate expression of proteins including transcription factors that transcriptionally control expression of other proteins (23–25). Likewise, RAG DSBs generated in lymphoid progenitor cells cause heritable changes in gene transcription that enhance the differentiation and function of NK cells and other mature innate lymphoid cells (33). In normal mice, several rounds of Vα-to-Jα rearrangement typically occur on each allele in a DP thymocytes until formation of an αβ TCR that promotes positive selection (47). The decreased Vα-to-Jα rearrangements in DP thymocytes of *PG* mice would correspond with fewer RAG DSBs to signal transcriptional changes in the expression of proteins that govern selection and/or differentiation of αβ T lineage cells. One of these proteins could enhance Zap70 protein expression, while another might promote NKT1 cell differentiation. Alternatively, Rag1^P326G^ mutation could prevent Rag1 from recruiting and/or ubiquitylating proteins that activate signals from RAG DSBs to modulate the expression and/or activity of proteins including Zap70. We did not detect differences in Zap70 activation or other aspects of TCR signaling between *WT* and *PG* thymocytes after anti-CD3 stimulation (data not shown). However, this approach is not as sensitive for assaying signalling differences as using tetramers or DCs to present peptides to transgenic αβ TCRs of fixed specificity (9, 48, 49)., which will be necessary to determine if Rag1^P326G^ mutation diminishes TCR signaling in DP thymocytes. A positive outcome would warrant determining the contribution of reduced Zap70 versus other protein expression changes.

The requirement for TCR selection on self-peptides displayed by self-proteins presents major challenges for establishing conventional and non-conventional αβ T cell populations that recognize and respond to a vast array of diverse foreign peptides, but not self-antigens. Thus, cell intrinsic mechanisms have evolved to render thymocytes more sensitive to TCR stimulation than their mature αβ T cell counterparts (48, 50–53), a phenomenon called thymic TCR tuning (49, 54–58). Our data are consistent with the possibility that the RAG1 RING domain ubiquitin ligase enhances thymic TCR tuning. Regardless, our study provides a phenotypic framework from which to elucidate precise molecular mechanisms by which the RAG1 RING domain regulates TCR gene assembly and selection to establish replete populations of αβ T lineage cells and central tolerance.

## Materials and Methods

### Mice

C57BL/6 background *PG* mice were made through gene-targeting as performed by InGenious Targeting Laboratory (2200 Smithtown Avenue, Ronkonkoma, NY). The 9.5 kb genomic DNA used to construct the targeting vector was sub-cloned from a 129Svev BAC clone. The CC>GG double point mutation was shuttled from pJMJ029[P326G](15) into the targeting vector using unique restriction sites, then confirmed by sequencing. Linearized targeting vector was electroporated into BA1 (C57BL/6 x 129/SvEv) embryonic stem (ES) cells. After selection with G418 antibiotic, clones were expanded for PCR analysis to identify recombinant ES clones. Positive clones were screened for short arm integration and then confirmation of the point mutation was performed by PCR and sequencing. Positive clones were analyzed by Southern blot to confirm long arm integration. Injected blastocysts were grown in foster mothers and chimeras were backcrossed against C57BL/6 mice to generate heterozygotes. These were crossed with CMV-*cre* mice (JAX #006054)(60) to remove the *neo* reporter cassette and generate the *Rag1^P326G^* allele. After confirmation of *neo* cassette removal, the strain was backcrossed against C57BL/6, and removal of CMV-*cre* was confirmed by PCR. Sequencing of the entire *Rag1* locus in progeny confirmed the intended CC>GG mutation and an additional T>C point mutation that generates a V238A substitution outside of the RING finger domain. The strain was deposited at the Mutant Mouse Regional Resource Center at the University of North Caroline (RAG1-tm1, MMRRC #: 37105). Before initiating studies described herein, the *Rag1^P326G^* allele was backcrossed for nine generations onto the C57BL/6 background. C57BL/6 background wild-type and *Rag1^-/-^* mice and BALB/c background *Rag1^-/-^* mice were purchased from Jackson Laboratories. All experimental mice were littermates or age-matched with control wild-type animals. All experiments were conducted in accordance with national guidelines and approved by the Institutional Animal Care and Use Committee (IACUC) of the Children’s Hospital of Philadelphia.

### Flow Cytometry

Single cell suspensions of all organs were ACK lysed to remove erythrocytes before being stained with LIVE/DEAD™ Fixable Aqua Dead Cell Stain Kit (Invitrogen) and antibodies against respective surface antigens (BD Biosciences, eBioscience, and Biolegend). Cells were stained for respective intracellular proteins using Cytofix/Cytoperm Fixation/Permeabilization solution kit (BD Biosciences). For staining of intracellular cytokines, cells (1 x10^6^) were cultured in the absence or presence of 50 ng/ml PMA (Sigma) and 1μg/ml Ionomycin (Cell Signaling Technology), with 2 μg/ml brefeldin A (Sigma) and 2 μM monensin (eBioscience) for 4 hours at 37°C. After staining for LIVE/DEAD and surface antigens, cells were stained for respective cytokines using the Cytofix/Cytoperm kit (BD Bioscience) or transcription factors using the FoxP3/Transcription Factor staining kit (eBiosciences). Data were acquired on a MACSQuant flow cytometer (Miltenyi Biotec) or LSRII Fortessa (BD Biosciences) and analyzed using Flowjo software version 10.5.3 (Tree Star). Analyses were conducted with the following antibodies: anti- CD4 (APC-eFluor 780, clone RM4-5, 1:100), anti-CD25 (BUV395, clone PC61, 1:100), anti-TCRβ (BV711, clone H57-597, 1:100), anti-CD19 (PE, clone 1D3, 1:100), anti-CD11b (PE, clone M1/70, 1:100), anti-CD11c (PE, clone HL3, 1:100), anti-TER (PE, clone TER-119, 1:100), anti-TCRγ/δ (PE, clone GL3, 1:100), anti-NK-1.1 (PE, clone PK136, 1:100), and anti-B220 (PE, clone RA3-6B2, 1:100) from BD Biosciences; or anti-CD8α (Pacific Blue, clone 53-6.7, 1:100), anti-CD44 (APC, PerCP/Cy5.5, clone IM7, 1:100), anti-CD24 (PE/Cy7, clone M1/69, 1:100), anti-CD28 (FITC, clone E18, 1:50), anti-CD45.1 (APC, clone A20, 1:100), anti-C45.2 (FITC, clone 104, 1:100) from BioLegend.

### Cell Sorting

To isolate DN3 cells, thymocytes were stained with PE-labelled CD4, CD8, CD11b, CD11c, NK1.1, Gr1, and Ter119. Non-labelled cells were enriched by MACS depletion using anti-PE microbeads and LS columns (Miltenyi). Enriched cells were then stained with CD4, CD8, CD44, CD25 and anti-lineage/stage markers (TCRβ, B220, CD19, CD11b, CD11c, TCR*δ*, NK1.1, Ter119) antibodies. DN3 thymocytes were sorted on Lin^-^CD4^-^CD8^-^CD44^-^CD25^+^ phenotype. To isolate DP cells, thymocytes were stained with CD4, CD8, and anti-lineage (B220, CD19, CD11b, CD11c, TCR*δ*, NK1.1, Ter119) antibodies. DP thymocytes were sorted on Lin^-^CD4^+^CD8^+^ phenotype. The cells of interest were isolated using a FACSAria Fusion sorter (BD Biosciences).

### Analyzing *Tcrb* and *Tcra* Rearrangements

Genomic DNA was extracted from DN3 or DP cells using the DNeasy Blood and Tissue kit (Qiagen). To measure *Tcrb* rearrangements in DN3 cells, a Taqman PCR assay was used to quantify Vβ-to-DJβ1.1 and Vβ-to-DJβ2.1 rearrangement levels with a primer specific for each Vβ paired and a probe, FAM or HEX, specific for Jβ1.1 or Jβ2.1, respectively. Taqman PCR was performed with conditions according to the manufacturer’s instructions (IDT DNA) on the ViiA 7 system (Applied Biosystems). PCR of CD19 was used for normalization. Primers, probes, and reaction conditions are as described (61). To assay *Tcrb* repertoire, DNA from DN3 thymocytes were sent to Adaptive Biotechnologies, who used multiplex PCR to amplify and deep sequence Vβ-Dβ-Jβ rearrangements. Gene segment usage was analyzed by ImmunoSEQ Analyzer software (Adaptive Biotechnologies). To assess *Tcra* rearrangements in DP cells, representative Vα-Jα rearrangements were quantified using a QuantiFast SYBR Green PCR kit (Qiagen) on the ViiA 7 system (Applied Biosystems) as described (62, 63). PCR of β2M was used for normalization. We quantified Vα14-Jα18 rearrangments as described (64).

### RT-qPCR for mRNA Expression

Respective cell populations were lysed in RLT buffer (Qiagen) containing 2-mercaptoethanol or TRIzol (Life Technologies) immediately after cell sorting. Total RNA was isolated using RNeasy Mini kit (Qiagen), treated with DNase (RNase-Free DNase Set, Qiagen), and reverse transcribed to generate cDNA with High-Capacity RNA-to-cDNA™ Kit (Themo-Fisher Scientific) according to manufacturer’s directions. The cDNA was subjected to qPCR using Power SYBR Green kit (Applied Biosystems) and actin, HPRT, ZAP70, and SYK primers (Qiagen). Relative expression was calculated using the ddCt method, using actin or HPRT as a housekeeping gene, and relevant calibrator sample (explained in figure legends for respective samples).

### Competitive Mixed Bone Marrow Chimeric Mice

6-week-old CD45.1 B6.SJL (WT) hosts were lethally irradiated (950 Rad on an X-RAD irradiator). Bone marrow from at least 3 pooled donors (CD45.1 WT only, CD45.2 RAG P326G mutant only, or mixed) was sterilely prepped and RBCs lysed. Acceptable mixing (CD45.1/CD45.2 ratio) was confirmed by flow cytometry. Then, 3-5×10^6^ total cells were injected intravenously into hosts 6 hours after irradiation. Mice were monitored over 8 weeks for graft rejection and graft-versus-host disease prior to euthanasia for analysis. Engraftment was measured using the following antibodies from Biolegend: anti-CD45.1 (APC- Cy7, clone A20, 1:100), anti-CD45.2 (FITC, Clone 104, 1:100), anti-NK1.1 (PE, clone PK136, 1:200), anti-CD44 (PerCP-Cy5.5, clone IM7, 1:100), anti CD24 (PE-Cy7, clone M1/69, 1:100), and anti-TCRβ (Pacific blue, clone H57-597, 1:100). CD1d tetramer from the NIH Tetramer Core (APC, 1:100) was a gift of Dr. Hamid Bassiri. Data were collected on a MACSQuant 10 (Miltenyi Biotec) and were analyzed using FlowJo software, version 10.

### *In vitro* T Cell Differentiation

Splenocytes from C57BL/6 background *Rag1^-/-^* mice were irradiated with 2,500 Rads (X-rad irradiator) and 3 x 10^5^ cells were seeded into each well of a 96 well plate. Naïve T cells (CD90.2^+^, CD4^+^, CD62L^+^, CD44^+^, CD25^-^) were sorted from C57BL/6 background *WT* or *Rag1^-/-^* mice using a FACSaria Fusion cell sorter (BD Biosciences), and 3 x 10^4^ sorted cells were added to each well containing irradiated feeder cells. All cells were cultured with 10 μg/ml anti-CD3 (Biolegend) and 3 μg/ml anti-CD28 (Biolegend) in RPMI with 10% FBS, 1% PSG, and 1x NEAA (Invitrogen), 1x Sodium Pyruvate (Invitrogen), and 0.001% 2-mercaptoethanol. rhIL-2 was used at 30 IU/mL, rmIL-12 and rmIL-4 at 10 ng/mL, rmIL-6 at 20 ng/mL, mTGFβ1 at 1 ng/mL, anti-mIL-4, anti-mIFNγ and anti-mIL-12 at 10 μg/mL (from Biolegend except rhIL- 2, rmIL-12, and rmIL-6 from Prepotech). The following cytokines/blocking antibodies were used to skew towards respective Th-subsets: Th0 condition: rhIL-2; Th1 condition: rhIL-2, rmIL-12 and anti-mIL-4; Th2 condition: rhIL-2, rmIL-4, anti-mIFNγ and anti-mIL-12; Th17 condition: rhIL-2, mIL-6, mTGFβ1, anti-mIL-4 and anti-mIL-12; Treg condition: rhIL-2, mTGFβ1, anti-mIL-4 and anti-mIL-12.

### LCMV Infections

Mice were infected intraperitoneally with 2 x 10^5^ plaque-forming units of LCMV-Armstrong and euthanized at indicated timepoints. Virus and gp33-tetramer were kindly provided by E. John Wherry (University of Pennsylvania).

### T Cell Transfer Model of Colitis

This model was performed as described (37). Briefly, naïve CD4^+^ T cells were sorted from respective donor mice (CD90.2^+^, CD4^+^, CD62L^+^, CD44^−^, CD25^−^) on a FACSAria Fusion cell sorter. 0.5 x 10^6^ cells were transferred in 100 μl of PBS into each of the donor mice via intraperitoneal injection. Non-transferred mice were injected with PBS. Mice were weighed weekly for the first 4 weeks, and then twice weekly for 4 weeks. Mice were euthanized 8 weeks post-transfer unless severe morbidity enforced premature euthanasia (in accordance with CHOP IACUC approved guidelines) or spontaneous death occurred.

### Statistical analysis

All data was analyzed in Graphpad Prism 8 using statistical tests indicated in the figure legends. Error bars indicate mean +/- SEM unless otherwise stated. n.s.=p>0.05, *p<0.05, **p<0.01, ***p<0.001, ****p<0.0001.

## Footnotes

This work was supported by the Fondation ARC pour la recherche sur le cancer (C.M.), La Ligue contre le cancer (C.M.), Fondation Pasteur Mutualite (C.M.), Nancy Taylor Foundation for Chronic Disease (E.M.B.) and NIH grants R03 AI097611 (J.M.J.), R01 HL112836 (E.M.B.), R01 AI121250-A1 (E.M.B.), R21 A1 135435 (C.H.B.), and RO1 AI 112621 (C.H.B.).

## Acknowledgements

The authors have no competing financial interests to declare.

---

**Supplemental Figure 1.**
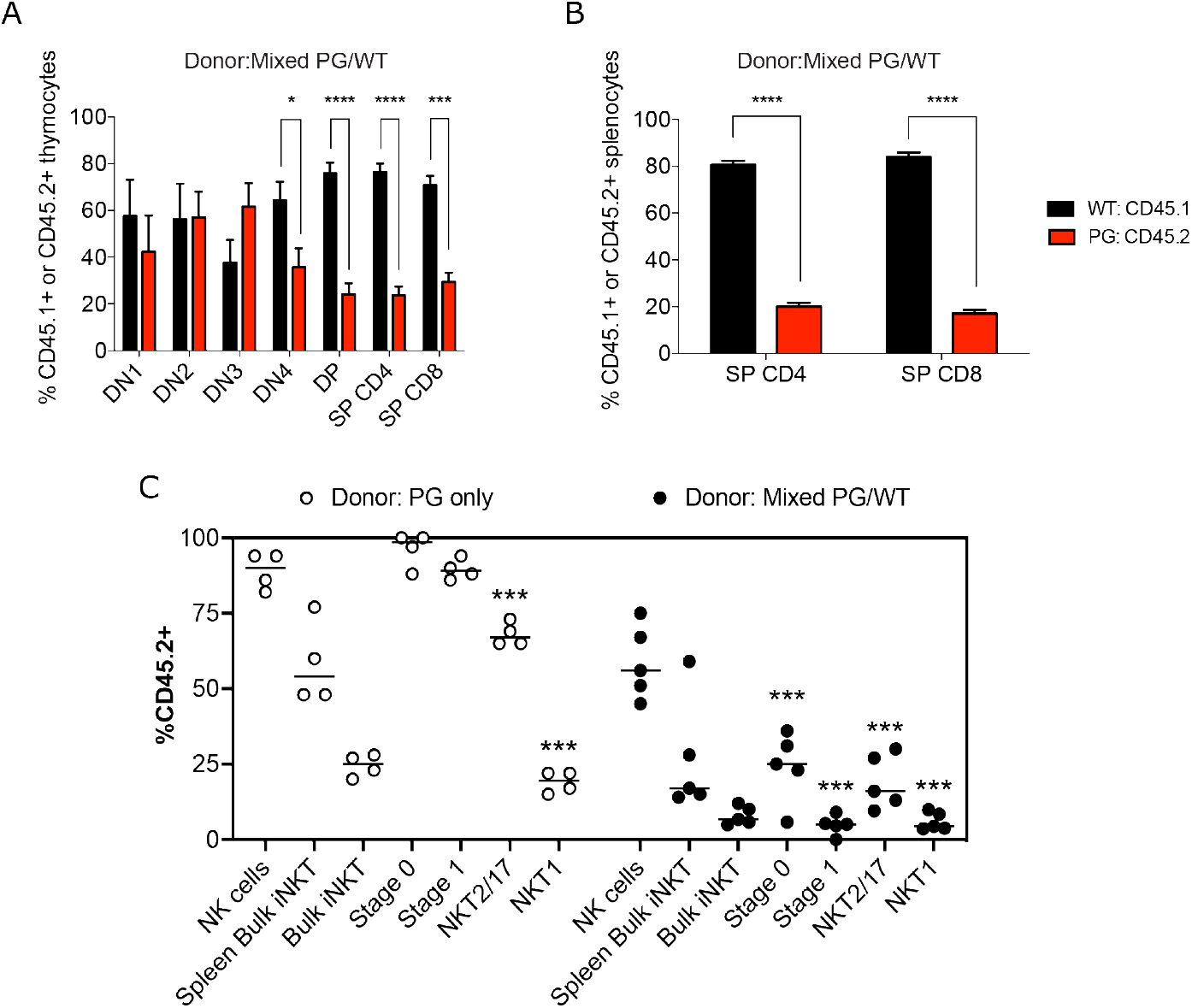
*PG* thymocytes exbibit cell intrinsic deficiencies in developmental progression. **(A-B)** Shown are percent of donor *PG* (CD45.2^+^) or *WT* (CD45.1^+^) cells within indicated (A) thymocyte or (B) splenocyte populations after transfer of a 1:1 mix of *PG* and *WT* bone marrow into lethally irradiated *WT* congenic (CD45.1^+^) recipients. Bone marrow was pooled from three donors of each genotype and injected into five recipients, all analyzed on the same day. Data represented as mean +/- SEM. Statistics calculated with multiple t tests using the Holm-Sidak method. *p<0.05, ***p<0.0005, ****p<0.00005. **(C)** Shown are percent donor *PG* (CD45.2^+^) chimerism in indicated compartments after lethal irradiation of *WT* congenic (CD45.1^+^) recipients. Open circles show percent *PG* chimerism after transfer of only *PG* bone marrow, while filled circles show *PG* chimerism in a 1:1 competitive setting with *WT* cells. *PG* bone marrow reconstitutes the blood NK cell compartment at expected ratios. In mice receiving only *PG* bone marrow, *PG* chimerism in bulk splenic iNKT (p<0.0001), bulk thymic iNKT (p<0.0001), NKT2/17 (p=0.001), and NKT1 (p<0.0001) are all significantly reduced relative to blood NK cells by one-way ANOVA with Dunnett multiple comparisons test. In mice receiving 1:1 mixed *PG* and *WT* bone marrow, *PG* chimerism in bulk splenic iNKT (p=0.0002), bulk thymic iNKT (p<0.0001), Stage 0 iNKT (p<0.0001), Stage 1 iNKT (p<0.0001), Stage 2 iNKT (p=0.001), and Stage 3 iNKT (p<0.0001) are all significantly reduced relative to blood NK cells. Bone marrow was pooled from three donors and injected into 4-5 mice per group, all analyzed on the same day.

**Supplemental Figure 2.**
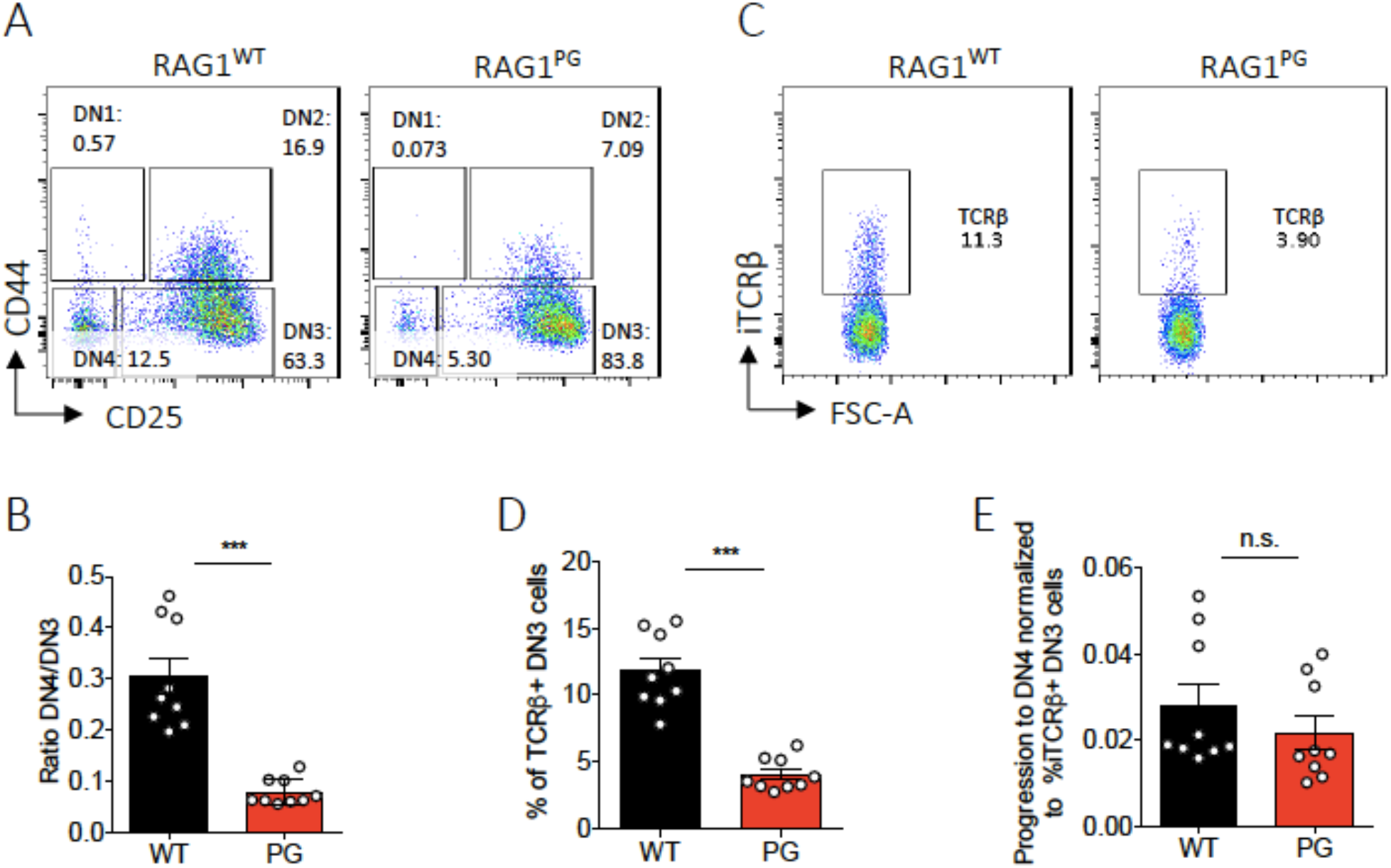
*PG* mice exhibit normal efficiency of pre-TCR signaled DN3-to-DN4 thymocyte differentiation. **(A)** Representative flow cytometry plots of DN stages of thymocyte development showing the frequency of cells at the DN3 and DN4 stages. **(B)** Quantification of percentages of DN3 cells that transition into DN4 cells. Shown are all data points and average values +/- SEM from three independent experiments, each with three mice of each genotype (n = 9 mice per genotype, analyzed by multiple t tests using the Holm-Sidak method; *** p<0.0005). **(C)** Representative flow cytometry plots of intracellular TCRβ protein expression in DN3 thymocytes showing the frequency of TCRβ^+^ cells in each gate. **(D)** Quantification of percentages of DN3 cells expressing intracellular TCRβ protein. **(E)** Ratios comparing the percentages of DN3 cells expressing intracellular TCRβ protein to the percentages of DN3 cells that transition into DN4 cells. (D-E) Shown are all data points and average values +/- SEM from three independent experiments each with three mice of each genotype (n = 9 mice per genotype, analyzed by unpaired t-test with Welch’s correction; n.s. p>0.05, *** p<0.0005

**Supplemental Figure 3.**
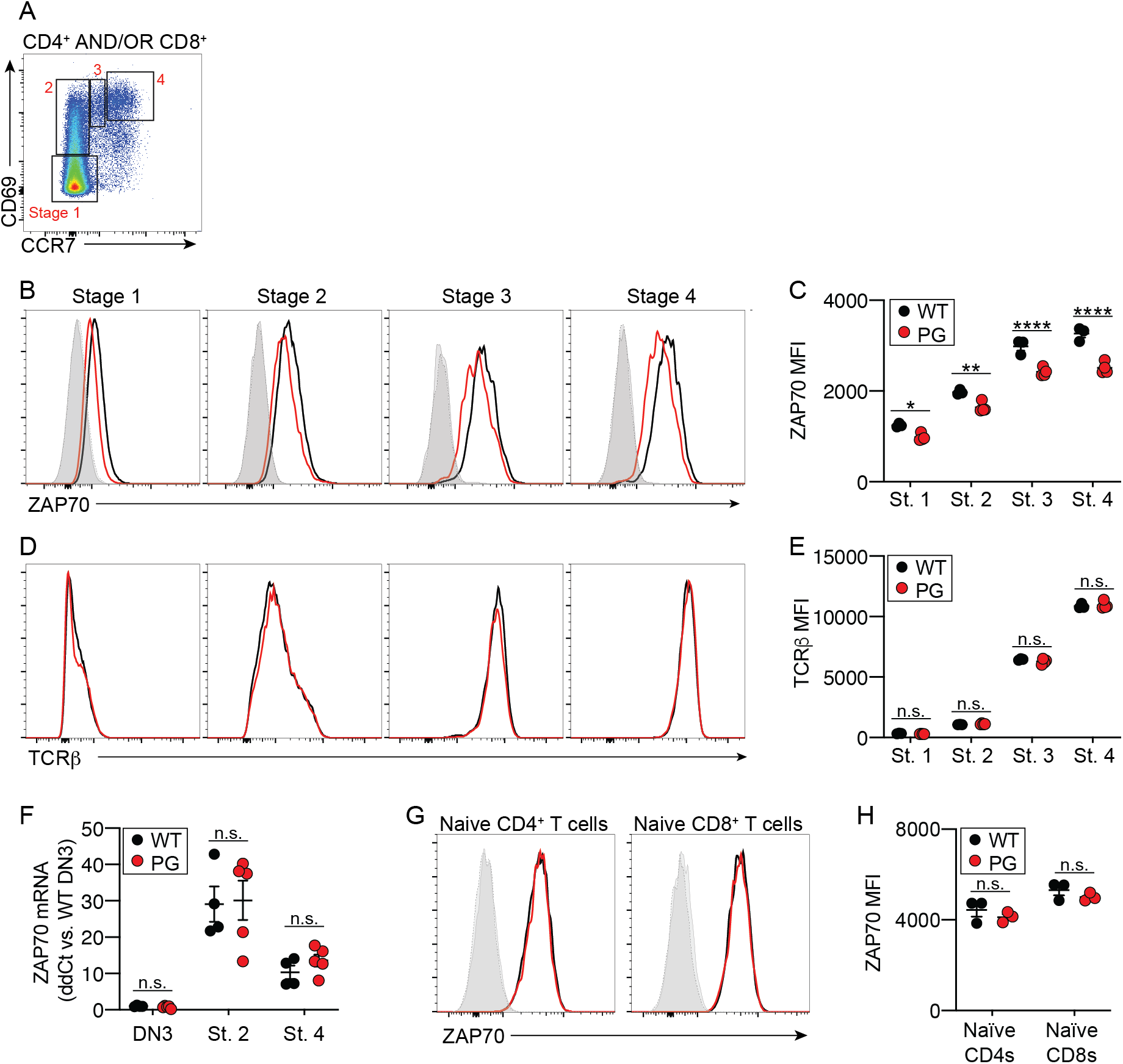
Thymocytes of PD mice have reduced Zap70 protein levels **(A)** Example gating strategy for Stage 1 - Stage 4 selecting thymocytes as depicted in Figure 4 and 5. Pre-gating: Live, singlets, CD4^+^ AND/OR CD8^+^. Stage 1 = CD69^-^CCR7^-^, Stage 2 = CD69^+^CCR7^-^, Stage 3 = CD69^+^CCR7^int^, Stage 4 = CD69^+^CCR7^hi^. **(B-E)** Representative flow cytometry histogram plots (B, D) and quantification (C, E) of Zap70 (B, C) and TCRβ (D, E) protein in Stages 1-4 of αβ TCR selection. (C, E) Shown are all data points and average values +/- SEM from one of three independent experiments (n = 4-5 mice per genotype, 2- way ANOVA with Sidak’s multiple comparison post-test) **(F)** qRT-PCR quantification of Zap70 mRNA in DN3 cells or Stage 2 or 4 cells. Relative mRNA levels were calculated using ddCt method with signals from each assay normalized to values from an assay for the Hprt gene. Shown are all data points and average values +/- SEM from one experiment (n = 4-5 mice per genotype, 2-way ANOVA with sidak multiple comparison post-test. **(G-H)** Representative flow cytometry histogram plots (G) and quantification (H) of Zap70 protein in naïve CD4^+^ or CD8^+^ T cells. Shown are all data points and average values +/- SEM from two independent experiments (n = 3 mice per genotype, 2-way ANOVA with Sidak multiple comparison post-test). For all graphs in figure: n.s. p>0.05; * p<0.05, **p<0.01, *** p<0.001, **** p<0.0001.

## Notes

### Competing Interest Statement

The authors have declared no competing interest.

## References

1. Bassing, C. H., W. Swat, and F. W. Alt. 2002. The mechanism and regulation of chromosomal V(D)J recombination. Cell 109 Suppl: S45–55.

2. Schatz, D. G., and P. C. Swanson. 2011. V(D)J recombination: mechanisms of initiation. Annu Rev Genet 45: 167–202.

3. Bhandoola, A., H. von Boehmer, H. T. Petrie, and J. C. Zuniga-Pflucker. 2007. Commitment and developmental potential of extrathymic and intrathymic T cell precursors: plenty to choose from. Immunity 26: 678–689.

4. von Boehmer, H., and F. Melchers. 2010. Checkpoints in lymphocyte development and autoimmune disease. Nat Immunol 11: 14–20.

5. Carico, Z., and M. S. Krangel. 2015. Chromatin Dynamics and the Development of the TCRalpha and TCRdelta Repertoires. Adv Immunol 128: 307–361.

6. Hogquist, K. A., and S. C. Jameson. 2014. The self-obsession of T cells: how TCR signaling thresholds affect fate ’decisions’ and effector function. Nat Immunol 15: 815–823.

7. Gascoigne, N. R., V. Rybakin, O. Acuto, and J. Brzostek. 2016. TCR Signal Strength and T Cell Development. Annu Rev Cell Dev Biol 32: 327–348.

8. Klein, L., B. Kyewski, P. M. Allen, and K. A. Hogquist. 2014. Positive and negative selection of the T cell repertoire: what thymocytes see (and don’t see). Nat Rev Immunol 14: 377–391.

9. Fu, G., V. Rybakin, J. Brzostek, W. Paster, O. Acuto, and N. R. Gascoigne. 2014. Fine-tuning T cell receptor signaling to control T cell development. Trends Immunol 35: 311–318.

10. Jones, J. M., and M. Gellert. 2003. Autoubiquitylation of the V(D)J recombinase protein RAG1. Proc Natl Acad Sci U S A 100: 15446–15451.

11. Singh, S. K., and M. Gellert. 2015. Role of RAG1 autoubiquitination in V(D)J recombination. Proc Natl Acad Sci U S A 112: 8579–8583.

12. Jones, J. M., A. Bhattacharyya, C. Simkus, B. Vallieres, T. D. Veenstra, and M. Zhou. 2011. The RAG1 V(D)J recombinase/ubiquitin ligase promotes ubiquitylation of acetylated, phosphorylated histone 3.3. Immunol Lett 136: 156–162.

13. Grazini, U., F. Zanardi, E. Citterio, S. Casola, C. R. Goding, and F. McBlane. 2010. The RING domain of RAG1 ubiquitylates histone H3: a novel activity in chromatin-mediated regulation of V(D)J joining. Mol Cell 37: 282–293.

14. Deng, Z., H. Liu, and X. Liu. 2015. RAG1-mediated ubiquitylation of histone H3 is required for chromosomal V(D)J recombination. Cell Res 25: 181–192.

15. Simkus, C., P. Anand, A. Bhattacharyya, and J. M. Jones. 2007. Biochemical and folding defects in a RAG1 variant associated with Omenn syndrome. J Immunol 179: 8332–8340.

16. Villa, A., C. Sobacchi, L. D. Notarangelo, F. Bozzi, M. Abinun, T. G. Abrahamsen, P. D. Arkwright, M. Baniyash, E. G. Brooks, M. E. Conley, P. Cortes, M. Duse, A. Fasth, A. M. Filipovich, A. J. Infante, A. Jones, E. Mazzolari, S. M. Muller, S. Pasic, G. Rechavi, M. G. Sacco, S. Santagata, M. L. Schroeder, R. Seger, D. Strina, A. Ugazio, J. Valiaho, M. Vihinen, L. B. Vogler, H. Ochs, P. Vezzoni, W. Friedrich, and K. Schwarz. 2001. V(D)J recombination defects in lymphocytes due to RAG mutations: severe immunodeficiency with a spectrum of clinical presentations. Blood 97: 81–88.

17. Zheng, N., P. Wang, P. D. Jeffrey, and N. P. Pavletich. 2000. Structure of a c-Cbl-UbcH7 complex: RING domain function in ubiquitin-protein ligases. Cell 102: 533–539.

18. Albert, T. K., H. Hanzawa, Y. I. Legtenberg, M. J. de Ruwe, F. A. van den Heuvel, M. A. Collart, R. Boelens, and H. T. Timmers. 2002. Identification of a ubiquitin-protein ligase subunit within the CCR4-NOT transcription repressor complex. EMBO J 21: 355–364.

19. Beilinson, H. A., R. A. Glynn, A. D. Yadavalli, J. Xiao, E. Corbett, H. Saribasak, R. Arya, C. Miot, Bhattacharyya J. M., Jones, J. M. R. Pongubala, C. H. Bassing, and D. G. Schatz. 2021. The RAG1 N-terminal region regulates the efficiency and pathways of synapsis for V(D)J recombination. J Exp Med 218.

20. Liang, H. E., L. Y. Hsu, D. Cado, L. G. Cowell, G. Kelsoe, and M. S. Schlissel. 2002. The “dispensable” portion of RAG2 is necessary for efficient V-to-DJ rearrangement during B and T cell development. Immunity 17: 639–651.

21. Horowitz, J. E., and C. H. Bassing. 2014. Noncore RAG1 regions promote Vbeta rearrangements and alphabeta T cell development by overcoming inherent inefficiency of Vbeta recombination signal sequences. J Immunol 192: 1609–1619.

22. Williams, J. A., K. S. Hathcock, D. Klug, Y. Harada, B. Choudhury, J. P. Allison, R. Abe, and R. J. Hodes. 2005. Regulated costimulation in the thymus is critical for T cell development: dysregulated CD28 costimulation can bypass the pre-TCR checkpoint. J Immunol 175: 4199–4207.

23. Bredemeyer, A. L., B. A. Helmink, C. L. Innes, B. Calderon, L. M. McGinnis, G. K. Mahowald, E. J. Gapud, L. M. Walker, J. B. Collins, B. K. Weaver, L. Mandik-Nayak, R. D. Schreiber, P. M. Allen, M. J. May, R. S. Paules, C. H. Bassing, and B. P. Sleckman. 2008. DNA double-strand breaks activate a multi-functional genetic program in developing lymphocytes. Nature 456: 819–823.

24. Bednarski, J. J., A. Nickless, D. Bhattacharya, R. H. Amin, M. S. Schlissel, and B. P. Sleckman. 2012. RAG-induced DNA double-strand breaks signal through Pim2 to promote pre-B cell survival and limit proliferation. J Exp Med 209: 11–17.

25. Bednarski, J. J., R. Pandey, E. Schulte, L. S. White, B. R. Chen, G. J. Sandoval, M. Kohyama, M. Haldar, A. Nickless, A. Trott, G. Cheng, K. M. Murphy, C. H. Bassing, J. E. Payton, and B. P. Sleckman. 2016. RAG-mediated DNA double-strand breaks activate a cell type-specific checkpoint to inhibit pre-B cell receptor signals. J Exp Med 213: 209–223.

26. Jordan, M. S., J. E. Smith, J. C. Burns, J. E. Austin, K. E. Nichols, A. C. Aschenbrenner, and G. A. Koretzky. 2008. Complementation in trans of altered thymocyte development in mice expressing mutant forms of the adaptor molecule SLP76. Immunity 28: 359–369.

27. Surh, C. D., and J. Sprent. 1994. T-cell apoptosis detected in situ during positive and negative selection in the thymus. Nature 372: 100–103.

28. Wilson, A., C. Marechal, and H. R. MacDonald. 2001. Biased V beta usage in immature thymocytes is independent of DJ beta proximity and pT alpha pairing. J Immunol 166: 51–57.

29. Rossjohn, J., D. G. Pellicci, O. Patel, L. Gapin, and D. I. Godfrey. 2012. Recognition of CD1d- restricted antigens by natural killer T cells. Nat Rev Immunol 12: 845–857.

30. Krovi, S. H., and L. Gapin. 2018. Invariant Natural Killer T Cell Subsets-More Than Just Developmental Intermediates. Front Immunol 9: 1393.

31. Dashtsoodol, N., S. Bortoluzzi, and M. Schmidt-Supprian. 2019. T Cell Receptor Expression Timing and Signal Strength in the Functional Differentiation of Invariant Natural Killer T Cells. Front Immunol 10: 841.

32. Hogquist, K., and H. Georgiev. 2020. Recent advances in iNKT cell development. F1000Res 9.

33. Karo, J. M., D. G. Schatz, and J. C. Sun. 2014. The RAG recombinase dictates functional heterogeneity and cellular fitness in natural killer cells. Cell 159: 94–107.

34. Jiang, W., M. S. Anderson, R. Bronson, D. Mathis, and C. Benoist. 2005. Modifier loci condition autoimmunity provoked by Aire deficiency. J Exp Med 202: 805–815.

35. Jin, Y., A. Lee, J. H. Oh, H. W. Lee, and S. J. Ha. 2019. The R229Q mutation of Rag2 does not characterize severe immunodeficiency in mice. Sci Rep 9: 4415.

36. Mottet, C., H. H. Uhlig, and F. Powrie. 2003. Cutting edge: cure of colitis by CD4+CD25+ regulatory T cells. J Immunol 170: 3939–3943.

37. Ostanin, D. V., J. Bao, I. Koboziev, L. Gray, S. A. Robinson-Jackson, M. Kosloski-Davidson, V. H. Price, and M. B. Grisham. 2009. T cell transfer model of chronic colitis: concepts, considerations, and tricks of the trade. Am J Physiol Gastrointest Liver Physiol 296: G135–146.

38. Eri, R., M. A. McGuckin, and R. Wadley. 2012. T cell transfer model of colitis: a great tool to assess the contribution of T cells in chronic intestinal inflammation. Methods Mol Biol 844: 261–275.

39. Livak, F., D. B. Burtrum, L. Rowen, D. G. Schatz, and H. T. Petrie. 2000. Genetic modulation of T cell receptor gene segment usage during somatic recombination. J Exp Med 192: 1191–1196.

40. Bortoluzzi, S., N. Dashtsoodol, T. Engleitner, C. Drees, S. Helmrath, J. Mir, A. Toska, M. Flossdorf, R. Ollinger, M. Solovey, M. Colome-Tatche, B. Kalfaoglu, M. Ono, T. Buch, T. Ammon, R. Rad, and M. Schmidt-Supprian. 2021. Brief homogeneous TCR signals instruct common iNKT progenitors whose effector diversification is characterized by subsequent cytokine signaling. Immunity 54: 2497–2513 e2499.

41. Cameron, G., D. G. Pellicci, A. P. Uldrich, G. S. Besra, P. Illarionov, S. J. Williams, N. L. La Gruta, J. Rossjohn, and D. I. Godfrey. 2015. Antigen Specificity of Type I NKT Cells Is Governed by TCR beta-Chain Diversity. J Immunol 195: 4604–4614.

42. Negishi, I., N. Motoyama, K. Nakayama, K. Nakayama, S. Senju, S. Hatakeyama, Q. Zhang, A. C. Chan, and D. Y. Loh. 1995. Essential role for ZAP-70 in both positive and negative selection of thymocytes. Nature 376: 435–438.

43. Hsu, L. Y., Y. X. Tan, Z. Xiao, M. Malissen, and A. Weiss. 2009. A hypomorphic allele of ZAP- 70 reveals a distinct thymic threshold for autoimmune disease versus autoimmune reactivity. J Exp Med 206: 2527–2541.

44. Sakaguchi, N., T. Takahashi, H. Hata, T. Nomura, T. Tagami, S. Yamazaki, T. Sakihama, T. Matsutani, I. Negishi, S. Nakatsuru, and S. Sakaguchi. 2003. Altered thymic T-cell selection due to a mutation of the ZAP-70 gene causes autoimmune arthritis in mice. Nature 426: 454–460.

45. Zhao, M., M. N. D. Svensson, K. Venken, A. Chawla, S. Liang, I. Engel, P. Mydel, J. Day, D. Elewaut, N. Bottini, and M. Kronenberg. 2018. Altered thymic differentiation and modulation of arthritis by invariant NKT cells expressing mutant ZAP70. Nat Commun 9: 2627.

46. Tuttle, K. D., S. H. Krovi, J. Zhang, R. Bedel, L. Harmacek, L. K. Peterson, L. L. Dragone, A. Lefferts, C. Halluszczak, K. Riemondy, J. R. Hesselberth, A. Rao, B. P. O’Connor, P. Marrack, J. Scott-Browne, and L. Gapin. 2018. TCR signal strength controls thymic differentiation of iNKT cell subsets. Nat Commun 9: 2650.

47. Huang, C. Y., B. P. Sleckman, and O. Kanagawa. 2005. Revision of T cell receptor {alpha} chain genes is required for normal T lymphocyte development. Proc Natl Acad Sci U S A 102: 14356–14361.

48. Davey, G. M., S. L. Schober, B. T. Endrizzi, A. K. Dutcher, S. C. Jameson, and K. A. Hogquist. 1998. Preselection thymocytes are more sensitive to T cell receptor stimulation than mature T cells. J Exp Med 188: 1867–1874.

49. Fu, G., J. Casas, S. Rigaud, V. Rybakin, F. Lambolez, J. Brzostek, J. A. Hoerter, W. Paster, O. Acuto, H. Cheroutre, K. Sauer, and N. R. Gascoigne. 2013. Themis sets the signal threshold for positive and negative selection in T-cell development. Nature 504: 441–445.

50. Lucas, B., I. Stefanova, K. Yasutomo, N. Dautigny, and R. N. Germain. 1999. Divergent changes in the sensitivity of maturing T cells to structurally related ligands underlies formation of a useful T cell repertoire. Immunity 10: 367–376.

51. Yachi, P. P., C. Lotz, J. Ampudia, and N. R. Gascoigne. 2007. T cell activation enhancement by endogenous pMHC acts for both weak and strong agonists but varies with differentiation state. J Exp Med 204: 2747–2757.

52. Pircher, H., U. H. Rohrer, D. Moskophidis, R. M. Zinkernagel, and H. Hengartner. 1991. Lower receptor avidity required for thymic clonal deletion than for effector T-cell function. Nature 351: 482–485.

53. Sant’Angelo, D. B., and C. A. Janeway, Jr. 2002. Negative selection of thymocytes expressing the D10 TCR. Proc Natl Acad Sci U S A 99: 6931–6936.

54. Fu, G., S. Vallee, V. Rybakin, M. V. McGuire, J. Ampudia, C. Brockmeyer, M. Salek, P. R. Fallen, J. A. Hoerter, A. Munshi, Y. H. Huang, J. Hu, H. S. Fox, K. Sauer, O. Acuto, and N. R. Gascoigne. 2009. Themis controls thymocyte selection through regulation of T cell antigen receptor-mediated signaling. Nat Immunol 10: 848–856.

55. Lesourne, R., S. Uehara, J. Lee, K. D. Song, L. Li, J. Pinkhasov, Y. Zhang, N. P. Weng, K. F. Wildt, L. Wang, R. Bosselut, and P. E. Love. 2009. Themis, a T cell-specific protein important for late thymocyte development. Nat Immunol 10: 840–847.

56. Johnson, A. L., L. Aravind, N. Shulzhenko, A. Morgun, S. Y. Choi, T. L. Crockford, T. Lambe, H. Domaschenz, E. M. Kucharska, L. Zheng, C. G. Vinuesa, M. J. Lenardo, C. C. Goodnow, R. J. Cornall, and R. H. Schwartz. 2009. Themis is a member of a new metazoan gene family and is required for the completion of thymocyte positive selection. Nat Immunol 10: 831–839.

57. Wang, D., M. Zheng, L. Lei, J. Ji, Y. Yao, Y. Qiu, L. Ma, J. Lou, C. Ouyang, X. Zhang, Y. He, J. Chi, L. Wang, Y. Kuang, J. Wang, X. Cao, and L. Lu. 2012. Tespa1 is involved in late thymocyte development through the regulation of TCR-mediated signaling. Nat Immunol 13: 560–568.

58. Li, Q. J., J. Chau, P. J. Ebert, G. Sylvester, H. Min, G. Liu, R. Braich, M. Manoharan, J. Soutschek, P. Skare, L. O. Klein, M. M. Davis, and C. Z. Chen. 2007. miR-181a is an intrinsic modulator of T cell sensitivity and selection. Cell 129: 147–161.

59. Marrella, V., P. L. Poliani, A. Casati, F. Rucci, L. Frascoli, M. L. Gougeon, B. Lemercier, M. Bosticardo, M. Ravanini, M. Battaglia, M. G. Roncarolo, M. Cavazzana-Calvo, F. Facchetti, L. D. Notarangelo, P. Vezzoni, F. Grassi, and A. Villa. 2007. A hypomorphic R229Q Rag2 mouse mutant recapitulates human Omenn syndrome. J Clin Invest 117: 1260–1269.

60. Schwenk, F., U. Baron, and K. Rajewsky. 1995. A cre-transgenic mouse strain for the ubiquitous deletion of loxP-flanked gene segments including deletion in germ cells. Nucleic Acids Res 23: 5080–5081.

61. Gopalakrishnan, S., K. Majumder, A. Predeus, Y. Huang, O. I. Koues, J. Verma-Gaur, S. Loguercio, A. I. Su, A. J. Feeney, M. N. Artyomov, and E. M. Oltz. 2013. Unifying model for molecular determinants of the preselection Vbeta repertoire. Proc Natl Acad Sci U S A 110: E3206–3215.

62. Shih, H. Y., J. Verma-Gaur, A. Torkamani, A. J. Feeney, N. Galjart, and M. S. Krangel. 2012. Tcra gene recombination is supported by a Tcra enhancer- and CTCF-dependent chromatin hub. Proc Natl Acad Sci U S A 109: E3493–3502.

63. Chen, L., Z. Carico, H. Y. Shih, and M. S. Krangel. 2015. A discrete chromatin loop in the mouse Tcra-Tcrd locus shapes the TCRdelta and TCRalpha repertoires. Nat Immunol 16: 1085–1093.

64. Gapin, L., J. L. Matsuda, C. D. Surh, and M. Kronenberg. 2001. NKT cells derive from double-positive thymocytes that are positively selected by CD1d. Nat Immunol 2: 971–978.

